# Somatic structural variation targets neurodevelopmental genes and identifies *SHANK2* as a tumor suppressor in neuroblastoma

**DOI:** 10.1101/572248

**Authors:** Gonzalo Lopez, Karina L. Conkrite, Miriam Doepner, Komal S. Rathi, Apexa Modi, Zalman Vaksman, Lance M. Farra, Eric Hyson, Moataz Noureddine, Jun S. Wei, Malcolm A. Smith, Shahab Asgharzadeh, Robert C. Seeger, Javed Khan, Jaime Guidry Auvil, Daniela S. Gerhard, John M. Maris, Sharon J. Diskin

## Abstract

Neuroblastoma is a malignancy of the developing sympathetic nervous system that accounts for 12% of childhood cancer deaths. Like many childhood cancers, neuroblastoma exhibits a relative paucity of somatic single nucleotide variants (SNVs) and small insertions and deletions (indels) compared to adult cancers. Here, we assessed the contribution of somatic structural variation (SV) in neuroblastoma using a combination of whole genome sequencing (WGS; n=135) and single nucleotide polymorphism (SNP) genotyping (n=914) of matched tumor-normal pairs. Our study design allowed for orthogonal validation and replication across platforms. SV frequency, type, and localization varied significantly among high-risk tumors. *MYCN* non-amplified high-risk tumors harbored an increased SV burden overall, including a substantial excess of tandem-duplication events across the genome. Genes disrupted by SV breakpoints were enriched in neuronal lineages and autism spectrum disorder (ASD). The postsynaptic adapter protein-coding gene *SHANK2*, located on chromosome 11q13, was disrupted by SVs in 14% of *MYCN* non-amplified high-risk tumors based on WGS and 10% in the SNP array cohort. Expression of *SHANK2* was low across human-derived neuroblastoma cell lines and high-risk neuroblastoma tumors. Forced expression of *SHANK2* in neuroblastoma cell models resulted in significant growth inhibition (P=2.62×10^-2^ to 3.4×10^-5^) and accelerated neuronal differentiation following treatment with all-trans retinoic acid (P=3.08×10^-13^ to 2.38×10^-30^). These data further define the complex landscape of structural variation in neuroblastoma and suggest that events leading to deregulation of neurodevelopmental processes, such as inactivation of *SHANK2*, are key mediators of tumorigenesis in this childhood cancer.

## INTRODUCTION

Neuroblastoma is a cancer of the developing sympathetic nervous system that most commonly affects children under five years of age, with a median age at diagnosis of 17 months (Maris 2010). Approximately 50% of cases present with disseminated disease at the time of diagnosis. Despite intense multi-modal therapy, the survival rate for this high-risk subset remains less than 50% (Maris 2010). Recent whole genome and exome sequencing studies of neuroblastoma have revealed relatively few recurrent protein-coding somatic mutations including single nucleotide variations (SNVs) and small (<50b) insertion/deletions (indels) (Cheung et al. 2012; Molenaar et al. 2012; Pugh et al. 2013; Sausen et al. 2013). Large-scale structural variations (SVs) such as deletions, insertions, inversions, tandem duplications and translocations can arise from mutational processes that alter chromosome structure and evade innate mechanisms of maintaining genomic stability. These diverse SVs are often acquired somatically in cancer and can act as driver mutations(Yang et al. 2013).

Multiple approaches to detect SVs in large array and sequencing datasets have been applied to cancer (Alkan et al. 2011; Yang et al. 2013; Tubio 2015; Macintyre et al. 2016). First, methods to identify copy number variations (CNVs) from intensity data (log R ratios) have been applied to single nucleotide polymorphism (SNP) genotyping and comparative genomic hybridization (CGH) arrays. More recently, these approaches were adapted and applied to read-depth measures from next generation sequencing. Numerous segmentation algorithms exist for both array (Carter 2007) and sequence-based (Zhao et al. 2013) approaches, with the resulting CNV calls ranging in size from a few hundred base-pairs to whole chromosomal alterations. Importantly, these calls are dosage-sensitive, allowing for numerical quantification of amplifications and deletions.

Analysis of CNVs in neuroblastoma primary tumors and matched blood samples led to identification of recurrent somatically acquired CNVs. These include focal amplification *of MYCN*, gain of chromosome 17q, and deletion of chromosomes 1p and 11q. These events are associated with an undifferentiated phenotype, aggressive disease, and poor survival (Deyell and Attiyeh; Brodeur et al. 1984; Gilbert et al. 1984; Seeger et al. 1985; Gehring et al. 1995; Caron et al. 1996; Plantaz et al. 1997; Bown et al. 1999; Guo et al. 1999; Maris et al. 2001; Łastowska et al. 2002; Attiyeh et al. 2005; Michels et al. 2007). In addition, focal deletions in the *ATRX* chromatin remodeler gene (*ATRX*) result in deleterious loss of function (Cheung et al. 2012; Kurihara et al. 2014). *ATRX* is implicated in the alternative lengthening of telomeres (ALT) phenotype. Focal CNVs involving other tumor suppressor genes, such as *PTPRD* (Stallings et al. 2006), *ARID1A* and *ARID1B* (Sausen et al. 2013) have also been reported.

While analysis of somatic CNVs has been incredibly useful, next generation sequencing (NGS) approaches can profoundly expand our understanding of SVs in cancer (Macintyre et al. 2016). Alignment-based methods to identify SVs focus on reads and read-pairs discordantly aligned to the reference genome. As such, these alignment-based approaches do not rely on dosage quantification and do not quantify numerical changes of deletions and tandem-duplications. However, they provide essential information about inversions, translocations and transposable elements, which are elusive to CNV callers. Furthermore, read coverage-based and alignment-based approaches have often been combined together to improve accuracy (Qi and Zhao 2011; Zhang and Wu 2011; Jiang et al. 2012); these and other available methods have been systematically reviewed (Tattini et al. 2015).

Recent studies employing alignment-based detection of SVs from WGS of primary neuroblastomas revealed structural rearrangements as key oncogenic drivers. These SVs mediate enhancer hijacking or focal enhancer amplification, influencing telomere maintenance through activation of telomerase reverse transcriptase gene (*TERT*) (Peifer et al. 2015; Valentijn et al. 2015; Kawashima et al. 2016) or deregulating the *MYC* oncogene (Zimmerman et al. 2018). Despite the demonstrated importance of somatic CNVs and other SVs in neuroblastoma, studies systematically integrating CNV and alignment based approaches are lacking. Therefore, the global landscape and mechanisms of pathogenicity for many of these events remain poorly understood.

Here, we studied the role of somatic SVs in a large neuroblastoma cohort comprised of 997 distinct primary neuroblastoma tumor-normal pairs obtained at diagnosis. Specifically, we integrated whole genome sequencing (WGS) from 135 tumor-normal pairs and 914 single nucleotide polymorphism (SNP) arrays. Alternative approaches to SV detection were considered for both datasets, which overlapped in a subset of 52 cases. As such, this study allowed for cross-platform validation of SVs. We further explored the functional impact of SVs by integrating matched transcriptomic and gene fusion data from 153 RNA-sequencing samples and 247 HumanExon arrays comprising 361 distinct tumor samples. Finally, we performed *in vitro* studies to assess the functional relevance of *SHANK2*, a newly identified tumor suppressor gene disrupted by SVs. Altogether, the integration of multi-omic datasets together with patient clinical profiles and biological experimentation serves to greatly expand the mutational landscape of neuroblastoma.

## RESULTS

### Patient characteristics and multi-omic datasets for the study of structural variations

To establish the landscape of SVs in neuroblastoma, we first sequenced the genomes of 135 primary diagnostic tumors and matched normal (blood leukocyte) DNA pairs through the Therapeutically Applicable Research to Generate Effective Treatments (TARGET) initiative (https://ocg.cancer.gov/programs/target). Samples were obtained through the Children’s Oncology Group (COG) and included 106 patients with high-risk tumors (29 *MYCN*-amplified and 77 non-*MYCN*-amplified), 14 with intermediate-risk tumors and 15 with low-risk tumors (**Fig. 1a, Supplementary Tables 1 and 2**). Whole genome sequencing (WGS) was performed by Complete Genomics (Drmanac et al. 2010) to a median average depth of 76x (**Supplementary Fig. 1a**) and primary data was processed via the Complete Genomics pipeline version 2.0. This pipeline reports small somatic variants (SNVs, small indels, and substitutions)(Carnevali et al. 2012), larger (>200bp) SVs, and read-depth coverage across the genome used to infer copy number segmentation profiles (**Online methods**).

**Figure 1:**
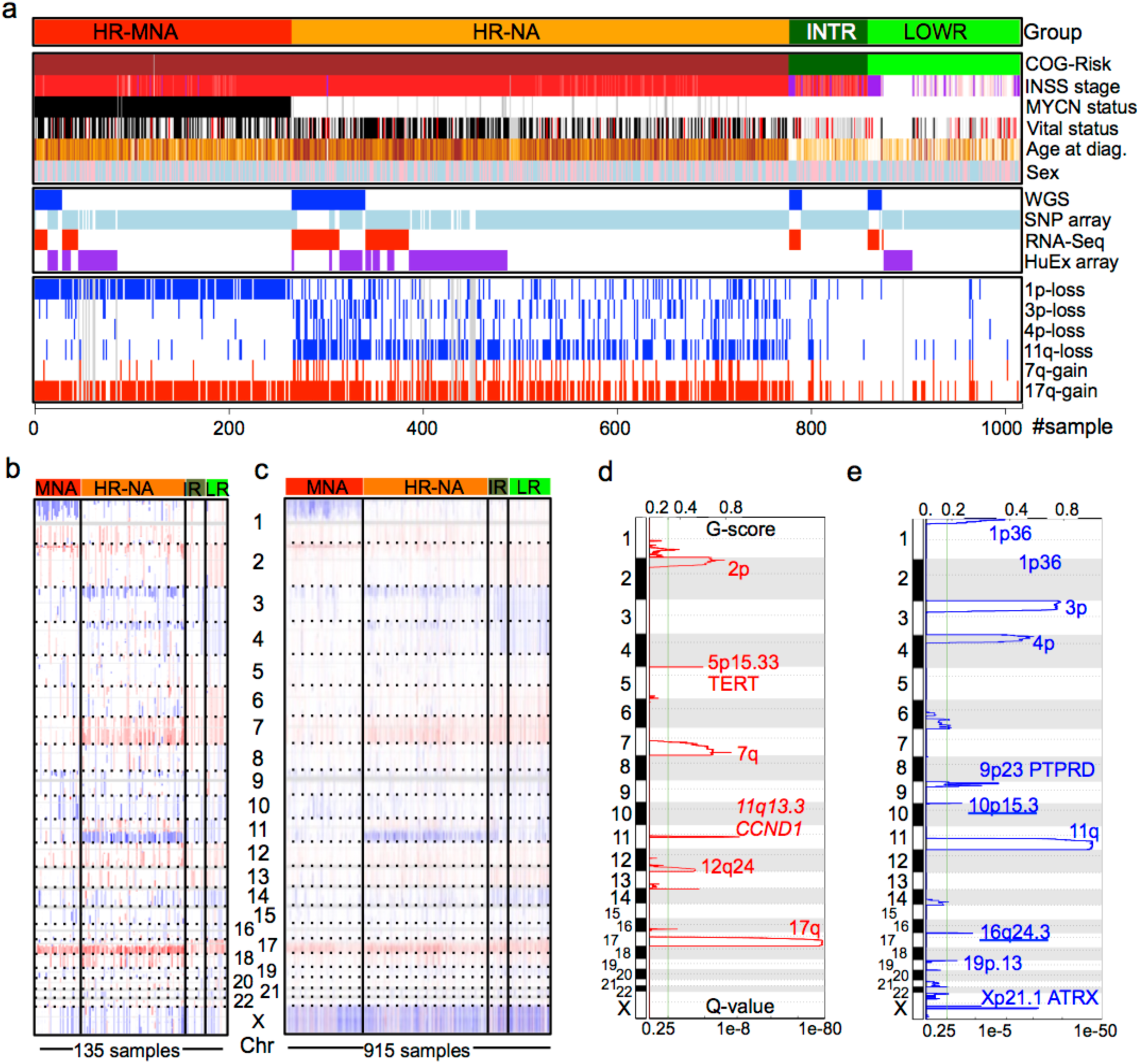
Novel somatic DNA copy number alterations (SCNAs) revealed by whole genome sequencing (WGS) of neuroblastoma tumors. **(a)** Survey of available samples, clinical information and data types used throughout this study (See also Supplementary Tables 1 & 2). **(b-c)** Integrated Genome Viewer (IGV) visualization of DNA copy number gains (red) and losses (blue) across neuroblastoma subtypes in the **(b)** WGS and **(c)** SNP datasets. **(d-e)** GISTIC q-value plots showing significant regions of **(d)** gain and **(e)** deletion in HR-NA samples in 77 samples derived from WGS dataset.

To augment the WGS data, and to provide independent replication, we genotyped and analyzed 914 patient tumor-normal pairs using Illumina SNP arrays (**Fig. 1a, Supplementary Tables 1 and 2**). This cohort included 696 high-risk (239 *MYCN*-amplified and 457 non-*MYCN*-amplified), 70 intermediate-risk and 145 low-risk tumor-blood pairs; 488 of these tumor samples were previously released (Attiyeh et al. 2009) and reanalyzed here. Copy number segmentation was obtained using the *SNPrank* algorithm implemented by the NEXUS^®^ software platform (**Online Methods**).

To further assess the biological relevance of SVs, we integrated additional data types generated through the TARGET initiative. These data included transcriptional profiles from RNA sequencing (N=153) and Affymetrix HumanExon arrays (HuEx, N=247). Patient clinical covariates were organized by the Children’s Oncology Group (COG) (**Fig. 1a, Supplementary Table 1 and 2;** https://ocg.cancer.gov/programs/target/data-matrix). Throughout the study, we examined disease risk groups as defined by the COG and the International Neuroblastoma Risk Group (INRG) (Cohn et al. 2009). Specifically, the following subtypes were considered: **LOWR**: low-risk neuroblastoma; **INTR:** intermediate-risk neuroblastoma; **MNA**: high-risk neuroblastomas with amplification of the *MYCN* oncogene, and **HR-NA**: high-risk neuroblastomas without *MYCN* amplification.

### Identification of novel regions of recurrent DNA copy number gain and loss

WGS-derived copy number profiles were compared with those obtained from the larger SNP array dataset. Somatic CNVs were visualized with Integrative Genome Viewer (IGV) and confirmed well-established patterns of large SCNAs in neuroblastoma that differed between the tumor clinical subtypes (**Fig. 1b, c**)(Wang et al. 2006; Michels et al. 2007). We further analyzed CNV segmentation profiles within neuroblastoma subtypes using GISTIC2.0 (Mermel et al. 2011). As expected, LOWR and INTR tumors harbored few focal and large CNVs, although aneuploidy was observed (**Supplementary Fig. 2a, b**). The MNA and HR-NA subsets shared highly recurrent 17q gains and *PTPRD* deletions (9p23) and differed in 2p24 gain (*MYCN* locus) and prevalence of deletions at 1p, 3p, 4p and 11q (**Fig. 1d, e, Supplementary Fig. 2c-e**). We also observed less frequently reported variants in HR-NA group, including deletions at 16q24.3 (Mosse et al. 2005) and segmental gains of the q-arm of chromosome 7, a region recently suggested to exhibit oncogenic potential in neuroblastoma (Bosse et al. 2017) (**Fig. 1d, Supplementary Fig. 2e**). CNV profiles derived from WGS are of higher resolution and returned significant peaks in the HR-NA subset not identified in SNP arrays. These CNVs included focal gains at chromosome 5p15.33 (Q-value=1.42 × 10^-3^) harboring the telomerase reverse transcriptase (*TERT*) gene) (**Fig. 1d**), intragenic deletions of the ATRX chromatin remodeler gene at Xq21.1, (Q-value=3.76 × 10^-3^), and a novel region of recurrent deletion at 10p15.3 (Q-value=6.16 × 10^-2^, **Fig. 1e**).

### Identification of SV breakpoints using orthogonal approaches: sequence junction (SJ-BP), read-depth (RD-BP) and copy number breakpoint (CN-BP) analyses

To strengthen our findings, we considered three approaches to SV breakpoint identification (**Table 1**). Specifically, we integrated alignment-based SV calls and read-depth CNVs from WGS as well as intensity-based CNV calls from genotyping arrays (**Table 1, Online Methods**). First, we obtained alignment-based SVs reported by the CGI somatic pipeline, which provides information about SV boundaries, size, and the type of variant in every sample. These included deletions (>500b), tandem-duplications (>40b), inversions (>30b), translocations, inversions and complex events (**Supplementary Fig. 1c-e**). We applied additional filters to remove likely artifacts including duplicate junctions across samples and common germline variants found in the Database of Genomic Variants (DGV; **Online Methods**) (MacDonald et al. 2014). This resulted in a total of 7,366 (**Supplementary Table 3**) SV calls distributed heterogeneously across neuroblastoma subtypes (**Fig. 2a**). These SVs were defined by sequence junctions delimited by two breakpoints in the genome, and will be referred to as sequence junction breakpoints (SJ-BP). We next mapped copy number dosage breakpoints derived from WGS read-depth segmentation profiles, hereafter referred to as read-depth breakpoints (RD-BP, **Online Methods**). A total of 2,836 RD-BPs were identified (µ=21) and were unevenly distributed across samples (**Fig. 2b**). Finally, analogous to the RD-BPs, we mapped copy number breakpoints from segmentation profiles derived from the larger SNP array cohort, referred to as copy number breakpoints (CN-BP, **Online Methods**). A total of 6,241 CN-BPs were identified across 914 samples (µ=6.8) (**Fig. 2c**). We subsequently verified the extent to which SV breakpoints overlapped between alternative methods and across WGS and SNP datasets. Overall, we observed high concordance, in agreement with benchmarks from other available methods (Qi and Zhao 2011; Zhang and Wu 2011; Jiang et al. 2012) (**Supplementary Fig. 3, Online Methods**). As expected from previous reports (Wang et al. 2006; Michels et al. 2007; Schleiermacher et al. 2012), we observed substantially more SV events in high-risk compared to intermediate- and low-risk tumors when considering SJ-BPs (**Fig. 2a**), RD-BPs (**Fig. 2b**) and CN-BP (**Fig. 2c**).

**Table 1.**
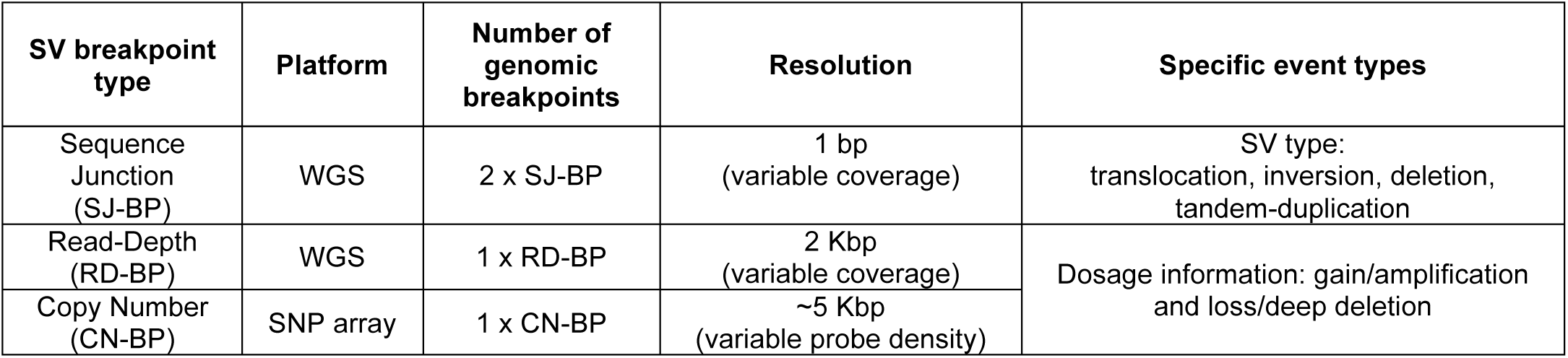
Definition of structural variant (SV) breakpoint analysis types.

**Figure 2:**
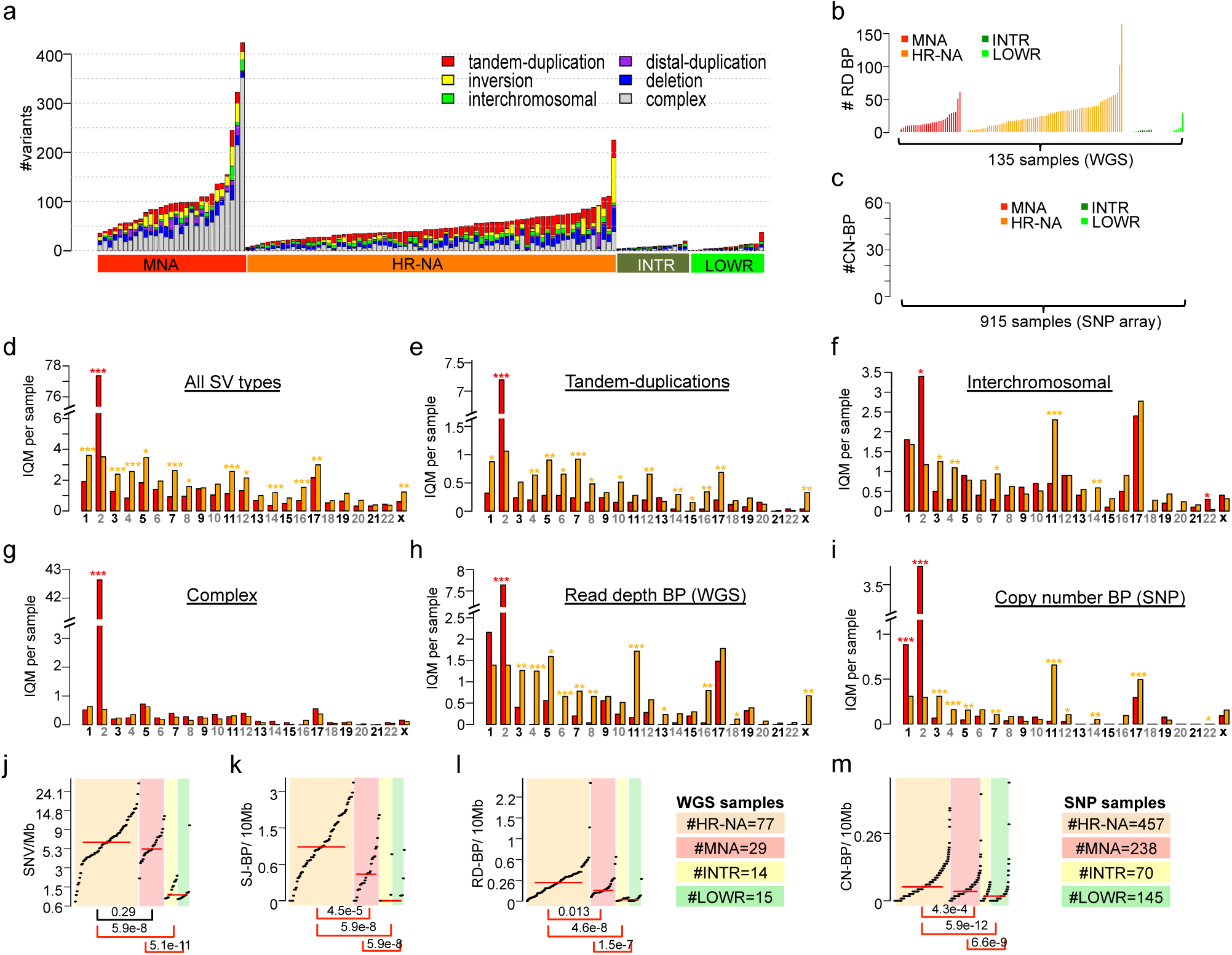
Somatic structural variation burden differs among neuroblastoma subtypes by quantity, type and genomic location. (a) Stacked bar chart of alignment based SV calls by type and neuroblastoma subtype in WGS dataset. (b) Bar plot representing the number of read-depth breakpoint (RD-BP) per sample across subtypes in the WGS dataset. (c) Bar plot representing the number of copy number breakpoint (CN-BP) per sample across subtypes in the SNP dataset. (d) Co-localization of RD-BPs with SJ-BP across WGS dataset samples and overall co-localization percentage (right bar). (e) Co-localization of SJ-BPs with RD-BP across WGS dataset samples and overall co-localization percentage (right bar). (f-k) By chromosome comparison between MNA and HR-NA of the inter-quantile average number of SVs including all SJ-BP variant types (f), duplications (g), inter-chromosomal (h), complex (i) as well as RD-BP and CN-BP. A Wilcoxon test is obtained for every chromosome and the p-value significance level is represented by asterisk (*** = p < 0.001, ** = P < 0.01, * = P < 0.05; asterisk color indicates the group with higher IQM, red=MNA and orange=HR-NA). Mutation burden analysis plot across neuroblastoma subtypes representing the burden of SNVs (l), SJ-BPs (m), RD-BPs (n) and CN-BP (o).

### Patterns of SV mutational burden differ across neuroblastoma high-risk subtypes

Comparison of SJ-BPs in MNA and HR-NA tumors revealed these high-risk subsets differed in SV type and genomic location (**Fig. 2d-g, Supplementary Fig. 4a-d**). MNA tumors harbored more SVs on chromosome 2 (P=1.6 × 10^-14^; **Fig. 2d**); however, these were largely confined to complex junctions at the *MYCN* amplicon at chromosome 2p24 (**Supplementary Fig. 4a**). Nearly all chromosomes displayed a higher frequency of SVs in HR-NA than MNA (**Fig. 2d**). Specifically, HR-NA tumors harbored more tandem-duplications in all chromosomes except chromosome 2 (P=4.0 × 10^-12^), in particular, chr7 (P=3.91 × 10^-5^), chr5 (P=1.2 × 10^-3^) and chromosome 4 (P=1.3 × 10^-3^) (**Fig. 2e**). Inter-chromosomal events were also more frequent in HR-NA tumors and overlapped with regions of known SCNAs other than chr2, including chromosome 3p (P=1.8 × 10^-3^), chromosome 4p (P=9.1 × 10^-6^) and chromosome 11q (P=1.9 × 10^-8^), but not chromosome 1p and 17q (**Fig. 2f**). In contrast, complex events showed no overall differences between high-risk groups with the exception of the aforementioned chr2 (**Fig. 2g**). Finally, RD-BP and CN-BP frequencies followed a similar pattern across chromosomes as that of SJ-BPs; MNA tumors harbored increased number of breakpoints in chromosomes 2 (P_RD-BP_=2.4 × 10^-9^, **Fig. 2h**; P_CN-_ BP=4.2 × 10^-83^, **Fig. 2i**) while HR-NA harbored increased frequencies in most other chromosomes and in particular, chromosome 11 (P_RD-BP_=2.0 × 10^-8^, **Fig. 2h**; P_CN-BP_=4.0 × 10^-25^, **Fig. 2i**).

We next studied overall differences in mutational burden and chromosomal instability across subtypes; we posit that the densities of breakpoints (SJ-BP, RD-BP and CN-BP) throughout the genome represent a *bona fide* measure of chromosomal instability (CIN). We also obtained measures of somatic SNV density. In order to avoid skewing of results due to the *MYCN* amplicon in MNA samples and regions exhibiting chromothripsis, we implemented an SNV and SJ-BP tumor burden measure robust against outliers. To this end, the genome was divided into 41 sequence mapped chromosome arms and the density of SVs per Mb was measured. For each sample, the interquartile mean (IQM) was derived from the 41 arm measurements (**Supplementary Fig. 4e,f**). Similarly, we obtained IQM density measurements from RD-BP and CN-BP chromosomal burdens. As expected, LOWR and INTR tumors carried very low mutational burden (**Fig. 2j-m**)(Wang et al. 2006; Michels et al. 2007). We observed increased CIN (SJ-BP, RD-BP and CN-BP) in HR-NA compared to MNA (Wilcoxon rank test: P_SJ-BP_=4.5 × 10^-5^, **Fig. 2k**; P_RD-BP_=1.3 × 10^-2^, **Fig. 2l**; P_CN-BP_=4.6 × 10^-8^, **Fig. 2m**), similar to previous reports(Caren et al. 2010). In contrast, no difference was observed in the average SNV burden (Wilcoxon rank test: P=0.29, **Fig. 2j**). These data support the notion that small SNVs and SVs arise from different mutational processes. These results also confirm the observation that HR-NA tumors exhibit increased CIN (Wang et al. 2006; Caren et al. 2010), and expand this to include other classes of SVs, such as tandem duplication events identified by SJ-BP analysis.

### Chromothripsis associates with major neuroblastoma oncogenic mechanisms

Previous studies have reported chromothripsis to occur in up to 18% of high-risk neuroblastomas (Molenaar et al. 2012) and identified associations between chromothripsis and key neuroblastoma oncogenes *MYCN* and *TERT* (Peifer et al. 2015; Valentijn et al. 2015). We therefore sought to leverage our large dataset to further explore the oncogenic associations of chromothripsis in neuroblastoma. We first identified alterations of major neuroblastoma oncogenes (*MYCN, TERT* and *ALK*) in our WGS and SNP cohorts. Rearrangements near the *TERT locus* were confirmed in 23 HR-NA samples and 2 MNA from the WGS dataset as well as 15 cases (14 HR-NA and 1 MNA) from the SNP dataset, one sample (PAPUTN) was present in both datasets (**Supplementary Fig 5a**). *TERT* expression was increased in these samples as well as in MNA tumors in accordance with previous reports (**Supplementary Fig. 5b,c**) (Peifer et al. 2015; Valentijn et al. 2015). Eleven cases from the WGS set with available DNA were validated using Sanger sequencing (**Supplementary Fig. 6**). In addition, CN-BPs were identified near *TERT* in fifteen HR-NA samples from the SNP array dataset (**Supplementary Fig 5a**), highlighting the ability of SNP arrays to detect these events using the CN-BP approach implemented in this study. *MYCN* amplification was determined diagnostically by FISH experiments in 29 samples from the WGS dataset (**Supplementary Table 2**); IGV visualization of segmentation data of 7Mb region surrounding *MYCN* confirmed these records (**Supplementary Fig. 7a**). We also explored events affecting the *ALK* gene. Two out of four rearrangements found near *ALK* also involved *MYCN*, (**Supplementary Fig. 7b**); these events were validated via Sanger sequencing (**Supplementary Fig. 7c**).

Next, chromothripsis was characterized based on clustered somatic rearrangements and alternating copy number states in defined chromosome regions (Maher and Wilson 2012). We identified candidate chromothripsis events at chromosome arms with unusual high breakpoint densities (> 2σ above the average of each sample’s breakpoint burden distribution) and a minimum of 6 breakpoints (both SJ-BPs and RD-BPs) in 27 regions (**Online Methods, Supplementary Table. 4**) involving 20 distinct high-risk tumors (19%). Chromothripsis was observed on chromosome 2 in a total of eight samples (**Table 2; Supplementary Fig. 8**); those samples showed enrichment in tumors harboring *MYCN* amplification (MNA) (7/8 samples, Binomial test P =7.4 × 10^-4^). Among them, two samples (PARETE and PATESI) involved co-amplification of *ALK* with *MYCN* (**Supplementary Fig. 8**). In addition, nine tumors harbored shattered chromosome 5p with strong enrichment in samples with rearrangements near *TERT* (8/9, Binomial test P = 7.3 × 10^-5^) (**Table 2**; **Supplementary Fig. 9**). Two samples (PAPSRJ and PAPUTN, **Supplementary Fig. 8**) included inter-chromosomal events involving the *MYCN* and *TERT* gene loci and co-amplification of both oncogenes. Other chromosomes involved in chromothripsis events included chromosome 1, 10, 11 and X in a female sample (**Supplementary Fig. 10**). Chromothripsis in most cases (15/20) was localized to a single chromosome involving either the whole chromosome (i.e. PATBMM, **Supplementary Fig. 9**) or local regions (i.e. PATESI, **Supplementary Fig. 8**). Multiple chromosomes were involved in 5/20 (25%) of cases with chromothripsis. One sample (PARIRD) harbored an event involving chromosomes 2, 17 and 22, while PANRVJ involved large regions of chromosomes 1 and 2 (**Supplementary Fig. 8**).

**Table 2.** Incidence of chromothripsis by chromosome across 135 WGS samples and incidence of high breakpoint density by chromosome across 914 SNP array samples.

| Chromosome |  | chr1 | chr2 | chr3 | chr4 | chr5 | chr6 | chr7 | chr8 | chr10 | chr11 | chr12 | chr13 | chr15 | chr17 | chr18 | chr19 | chr22 | chrX |
| --- | --- | --- | --- | --- | --- | --- | --- | --- | --- | --- | --- | --- | --- | --- | --- | --- | --- | --- | --- |
| WGS | HR_MNA (29) | 0 | 7 | 0 | 1 | 2 | 0 | 0 | 0 | 0 | 0 | 0 | 0 | 0 | 1 | 0 | 0 | 1 | 0 |
|  | HR_NA (77) | 2 | 1 | 0 | 0 | 7 | 0 | 0 | 0 | 1 | 2 | 0 | 0 | 0 | 0 | 0 | 1 | 0 | 0 |
|  | TERT SV+(25) | 1 | 2 | 0 | 1 | 8 | 0 | 0 | 0 | 1 | 0 | 0 | 0 | 0 | 0 | 0 | 1 | 0 | 0 |
|  | TERT WT (81) | 1 | 6 | 0 | 0 | 1 | 0 | 0 | 0 | 0 | 2 | 0 | 0 | 0 | 1 | 0 | 0 | 1 | 0 |
|  | Male (65) | 1 | 7 | 0 | 1 | 4 | 0 | 0 | 0 | 1 | 1 | 0 | 0 | 0 | 1 | 0 | 1 | 1 | 0 |
|  | Female (41) | 1 | 1 | 0 | 0 | 4 | 0 | 0 | 0 | 0 | 1 | 0 | 0 | 0 | 0 | 0 | 0 | 0 | 0 |
| SNP | HR_MNA (239) | 2 | 46 | 1 | 0 | 1 | 1 | 1 | 1 | 0 | 2 | 1 | 0 | 1 | 1 | 1 | 1 | 0 | 1 |
|  | HR_NA (428) | 2 | 0 | 3 | 4 | 10 | 2 | 2 | 1 | 0 | 5 | 5 | 2 | 1 | 2 | 1 | 3 | 0 | 6 |
|  | TERT SV+(30) | 0 | 0 | 0 | 1 | 7 | 0 | 0 | 0 | 0 | 1 | 0 | 0 | 0 | 0 | 0 | 0 | 0 | 1 |
|  | TERT Unk (670) | 4 | 47 | 4 | 3 | 4 | 3 | 3 | 2 | 0 | 6 | 6 | 2 | 2 | 3 | 2 | 4 | 0 | 6 |
|  | Male (401) | 4 | 27 | 2 | 1 | 6 | 1 | 1 | 2 | 0 | 3 | 3 | 2 | 1 | 1 | 1 | 2 | 0 | 1 |
|  | Female (299) | 0 | 20 | 2 | 3 | 5 | 2 | 2 | 0 | 0 | 4 | 3 | 0 | 1 | 2 | 1 | 2 | 0 | 6 |

\* Squared cells indicates association with clinical groups (red: P binomial test $< 0.001$ ; green: P binomial test $< 0.05$ )

We next sought further confirmation of our results in the larger SNP array dataset. In the absence of sequence junction information, we focused on unusual high density (>2σ above average) of CN-BPs (**Table 2)**. We observed high-breakpoint density on chromosome 2 enriched in MNA samples (46/46, P ∼ 0). We also detected enrichment of high breakpoint density on chromosome 5 involving cases harboring rearrangements or CN-BPs near *TERT* (7/11, P = 3.01 × 10^-8^). In addition, high breakpoint density on chromosome X was enriched in female patients (6/7, P = 4.7 × 10^-2^), although no specific oncogenic associations were determined. Overall, SNP array analysis of high CN-BP density supported and replicated observations from the WGS analysis.

### Identification of genes recurrently altered by SVs in high-risk neuroblastomas

In order to identify genes affected by recurrent somatic SVs in neuroblastoma, we generalized the approach described previously for *TERT, MYCN* and *ALK* genes. SVs were assigned to different categories according to the inferred impact on the exonic sequence of known RefSeq genes (**Fig. 3a,b**). Sequence junction breakpoints (SJ-BPs) provide more detailed information, including the type of SV and the two genomic breakpoint locations involved. With this knowledge, we classified SVs into: a) “coding” SVs that modify the exonic sequence of known genes including whole gene copy number alterations (duplications and deletions, size up to 2Mb), or b) “non-coding” SVs that do not modify the exonic sequences but might have an impact on regulatory regions proximal to known genes (100Kb upstream and 25Kb downstream) or intronic regions (**Fig. 3a**). In contrast to SJ-BP junctions obtained from discordantly aligned mate read pairs, dosage-based breakpoints (RD-BP and CN-BP) cannot identify their counterpart location in the genome. Therefore, events such as translocations and inversions cannot be defined. However, read-depth and array intensity based copy number inform about dosage gains and losses. With this in mind, we assumed the impact as a) “coding”: breakpoints within the transcription start and end positions of known genes, or b) “non-coding”, breakpoints located on proximal upstream and downstream regions (**Fig. 3b**). In addition, we localized CNVs involving amplification (CN_WGS_ > 8; CN_SNP_>4.5, **Online Methods**) and deep deletions (CN_WGS_ < 0.5; CN_SNP_ < 0.9, **Online Methods**) (**Fig. 3b**).

**Figure 3:**
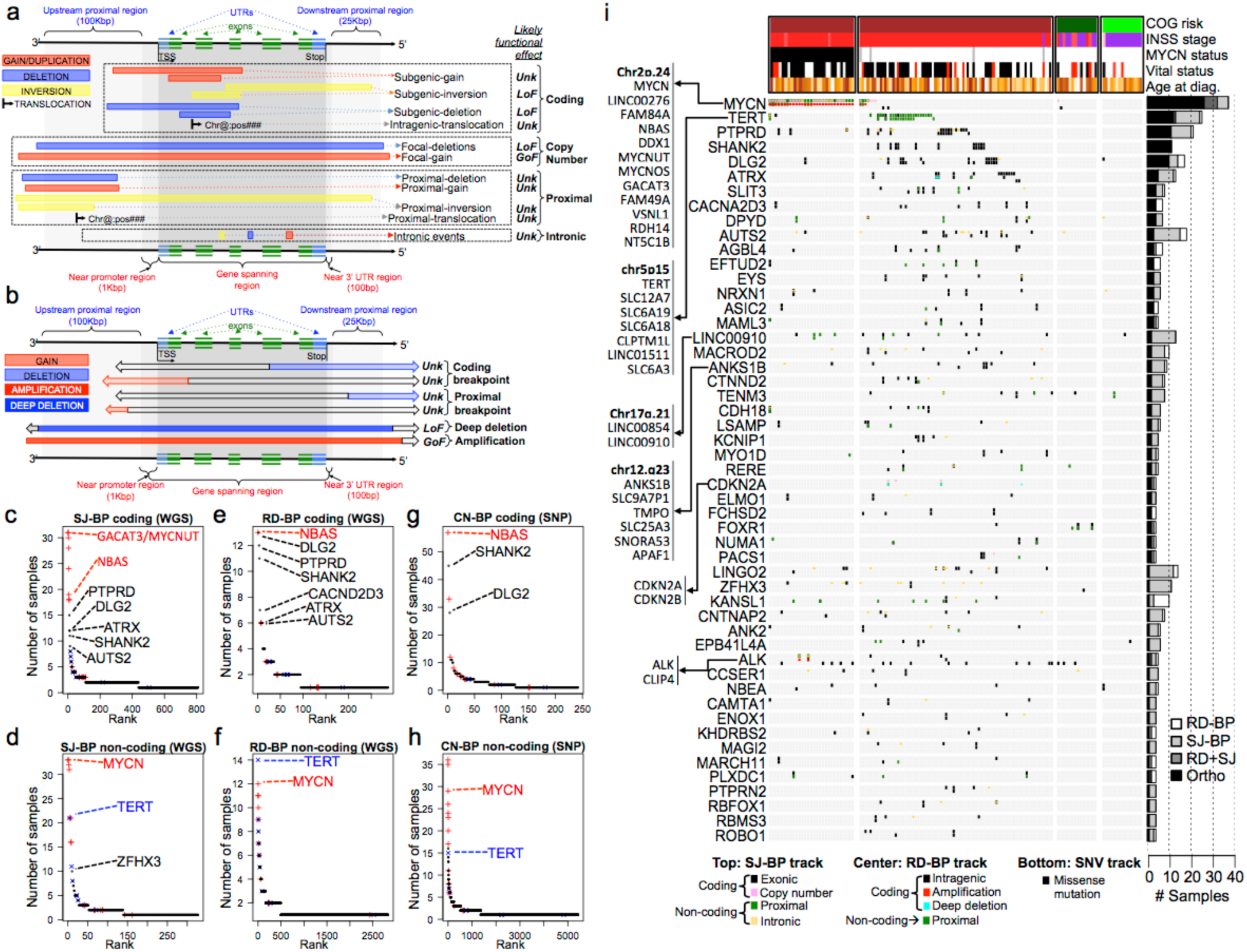
Identification of recurrently altered genes in neuroblastoma by breakpoint analyses. (a) Schematic representation of SVs derived from junction breakpoints (SJ-BPs)) classified according to their impact on known genes. (b) Schematic representation of read-depth and copy number breakpoints (RD-BPs, CN-BPs) classified according to their impact on known genes. (c-h) Recurrently altered genes rankings based on different breakpoint analyses and mode of impact: (c) gene coding sequences with recurrent SJ-BPs, (d) gene proximal and intronic sequences with recurrent SJ-BPs, (e) gene proximal sequences with recurrent RD-BPs (f) gene coding sequences with recurrent RD-BPs, (g) gene coding sequences with recurrent CN-BPs and (h) gene proximal sequences with recurrent CN-BPs. (i) Oncoprint based on the WGS dataset recurrently altered genes by SVs detected through orthogonal approaches (SJ-BP and RD-BP) as depicted in bar plot (right). The oncoprint aggregates three tracks per gene representing different BP analysis (upper =SJ-BP, center=RD-BP) and recurrent pathogenic SNVs (lower).

Based on the aforementioned definitions, we ranked recurrently altered genes according to the number of samples harboring “coding” and “non-coding” SVs for each of the 3 alternative breakpoint analyses (SJ-BPs, RD-BPs and CN-BPs; **Fig. 3c-h and Supplementary Table 5**). Overall, recurrently altered genes by ‘coding’ and ‘non-coding’ events return highly concordant results across the three approaches, with *MYCN, TERT*, and their neighbor genes at chr2.p24 and chr5.p15 (**Fig. 3c-g**) occupying top ranks. These were followed by known neuroblastoma altered genes *PTPRD* and *ATRX*. Notably, several top-ranked genes were novel, including *SHANK2* and *DLG2* located at chr11.q13 and chr11.q14 respectively, and others such as *AUTS2* at chr7.q11 and *CACNA2D3* at chr3.p14 (**Fig. 3c,e,g**). These analyses also reflect the long tail of altered genes of unknown pathogenicity (**Fig. 3c-g**).

In order to provide an integrated overview of the landscape of altered genes, we combined WGS based methods (SJ-BP and RD-BP) into a ranking of recurrently altered genes with co-localizing breakpoints, hence orthogonally validated. A total of 77 genes have at least 1 co-localizing SJ-BP and RD-BP breakpoint (**Fig. 3i, Supplementary Table 6**); in addition, we annotated likely pathogenic SNV calls (**Supplementary Table 7**). Many altered genes clustered in specific regions associated with known oncogenes such as chr2.p24 near *MYCN* (11 genes) and chr5.p15 near *TERT* (7 genes). The ranking was led by *MYCN* with 37 samples harboring variants. Orthogonal validation was obtained in 26 of these cases by co-localizing SJ-BP and RD-BP; these included 29 MNA and 8 HR-NA tumors. Interestingly, 11 HR-NA samples harbor alterations of *MYCN* (8 SVs and 3 SNVs) supporting the pathogenic role of *MYCN* in non-amplified tumors. *TERT* rearrangements were identified in 25 samples; orthogonal validation of breakpoints was observed in 12 cases (**Supplementary Fig. 5a**). *PTPRD* was altered in 20 samples, 11 of which were orthogonally validated (**Supplementary Fig. 11a**) (Stallings et al. 2006; Clark et al. 2012). We identified 12 *ATRX* intragenic deletions (5 orthogonally validated) and one tandem-duplication in HR-NA tumors (**Supplementary Fig. 11b**)(Cheung et al. 2012). Other genes with a known role in neuroblastoma and cancer were altered by SVs. Specifically, the *ALK* gene harbored SV rearrangements in five samples (**Supplementary Fig. 7b**). Another 18 samples exhibited somatic SNVs in *ALK*, resulting in a combined set of 23 samples being affected (17% of all neuroblastomas).

Genes lacking a well-established role in neuroblastoma were also disrupted by recurrent SVs in this study. Specifically, the *SHANK2* gene was disrupted in 11 HR-NA tumors; three samples involved gene fusions that did not appear in frame. *DLG2*, a newly described tumor suppressor in osteosarcoma (Smida et al. 2017; Shao et al. 2019), was disrupted in 10 samples based on SJ-BP and 14 samples based on RD-BP analyses, two of which involved gene fusion events. Both *SHANK2* and *DLG2* are located on chromosome 11q and play a role in the formation of postsynaptic density (PSD)(Kaizuka and Takumi 2018). Other novel candidate altered genes included *AUTS2* with frequent intragenic deletions at chr7q (N^orth^=3; N^tot^=18, **Supplementary Fig. 12a**) and the calcium channel *CACNA2D3* (N^orth^=4; N^tot^=11), which corresponded to a deletion breakpoint at chr3.p14.3 (**Supplementary Fig. 12b**). A region proximal to the *LINC00910* long non-coding RNA (lncRNA) on chromosome 17 suffered rearrangements in 13 tumors (**Supplementary Fig. 13a**). Finally, *CDKN2A* and *CDKN2B* focal deletions were identified in three tumors (**Supplementary Fig. 13b**).

### Sanger sequencing validation of SVs in high-risk neuroblastoma tumors

In addition to orthogonal validation by breakpoint analyses, we validated SV junctions by Sanger sequencing when samples had sufficient DNA available in our tumor bank (**Supplementary Table 8**). This validation effort focused on key genes and included twelve proximal *TERT* SVs (**Supplementary Fig. 6**), four *ATRX* deletions (**Supplementary Fig. 14**), four proximal *ALK* variants (**Supplementary Fig. 7c**), eleven *SHANK2* translocation events, nine of which involved chromosome 17q (**Supplementary Fig. 15**), and twelve *DLG2* variants (**Supplementary Fig. 16**). In total, we validated 45 SVs (**Supplementary Table 8**). The original CGI cancer pipeline classifies SVs into high and low confidence variants depending on the number of read pairs supporting the evidence (N_reads_ threshold = 10). Our pipeline rescued many cases classified as low-confidence by CGI. Specifically, six out of 45 SVs evaluated (13.3%, 3 *ATRX* and 3 *DLG2*) had low coverage (< 10 reads), but were confirmed by Sanger sequencing.

### SVs have a regional transcriptional effect in neuroblastoma tumors

To gain further understanding of the functional relevance of SVs, we performed an expression quantitative trait loci (eQTL) analysis for each of the recurrent SV-associated genes (**Supplementary Fig. 17a**). The analysis, which was replicated in the two available transcriptional datasets (RNA-seq and HuEx array), reported consistent up-regulation of *MYCN* and *TERT* including their neighbor genes in association with SVs. We also observed up-regulation of the lncRNA *LINC00910* (P_RNA_ = 7.0 × 10^-3^) at chr17.q21, a region with frequent inter-chromosomal translocations. In contrast, *CDKN2A* was down-regulated (P_HuEx_= 4.7 × 10^-2^; P_both_ = 2.5 × 10^-2^) by focal deletions and *PLXDC1* at chr17.q12 was down-regulated (P_HuEx_= 4.7 × 10^-2^; P_both_ = 2.5 × 10^-2^) in association with 17q gain breakpoints.

In addition to changes in overall gene expression by eQTL, translocations may lead to the expression of gene fusion transcripts. We explored RNA-seq samples with three available gene fusion methods (STAR-fusion(Dobin et al. 2013), fusionCATCHER(Daniel Nicorici 2014) and DeFUSE(McPherson et al. 2011) (**Supplementary Table 9**). The three methods return a wide range of fusion events (N_STAR-fusion_=24,837, N_fusionCATCHER_=6,898, N_DeFUSE_=22,837), and overlapped in only 68 events (0.1%, **Supplementary Fig 17b**). Notably, the subset of fusion events with matching translocation from WGS comprised 66 events (N_STAR-fusion_=45, N_fusionCATCHER_=36, N_DeFUSE_=44), with the three methods overlapping in 26 events (40%, **Supplementary Fig 17c**). These data show that DNA/RNA combined evidence verification returns high precision gene fusion events. The most frequent gene fusion event with both RNA and DNA evidence involved *SHANK2*; the three *SHANK2* fusion events involved 17q genes: *EFTUD2, MED1* and *FBXL20. DLG2* exhibited gene fusion events in two samples involving *SEMA6C* and *MYCBP2* at chromosomes 12 and 13 respectively. However, none of the *SHANK2* and *DLG2* fusion transcripts appeared to be in-frame, suggesting the fusion transcripts may not be biologically relevant and that these are more likely loss of function events. Conversely, we observed an in-frame fusion transcript and translocation involving *FOXR1:DDX6*, where oncogenic fusion events have previously been described in neuroblastoma (Santo et al. 2012).

### Neurodevelopmental genes are recurrently disrupted by SVs in neuroblastoma

In order to identify pathways targeted by SVs, we considered recurrently altered genes from each of the coding (N>2) and non-coding (N>3) altered gene lists (#genes: SJ-BP^coding^=109, SJ-BP^non-coding^=36, RD-BP^coding^=76, RD-BP^non-coding^=27, CN-BP^coding^=77 And CN-BP^non-coding^=88, **Fig. 3c-h**). We tested each gene list for enrichment across Gene Ontology, pathway, and disease gene classes using ToppGene (Chen et al. 2009) (**Supplementary Table 10**). Genes with coding sequences altered showed consistent results across the three breakpoint mappings, and revealed strong enrichment in genes involved in autism spectrum disorder (ASD) susceptibility (P_SJ-BP_ = 2.8 × 10^-9^; P_RD-BP_ = 2.9 × 10^-5^; P_CN-BP_= 2.7 × 10^-9^) and other neurodevelopmental disorders (NDD) as well as protein localization to synapse (P_SJ-BP_ = 1.2 × 10^-5^; P_RD-BP_ = 1.1 × 10^-7^; P_CN-BP_= 2.4 × 10^-6^) and other neuronal related classes (**Fig. 4a-c**; **Supplementary Table 10**). The gene sets with ‘non-coding’ alterations were more variable across the alternative breakpoint analyses, but were dominated by events involving *MYCN* and *TERT* in association with the disease class “stage, neuroblastoma” (P_SJ-BP_ = 1.9 × 10^-6^; P_RD-BP_ = 2.5 × 10^-5^; P_CN-BP_ = 9.2 × 10^-5^, **Supplementary Fig. 18**).

**Figure 4:**
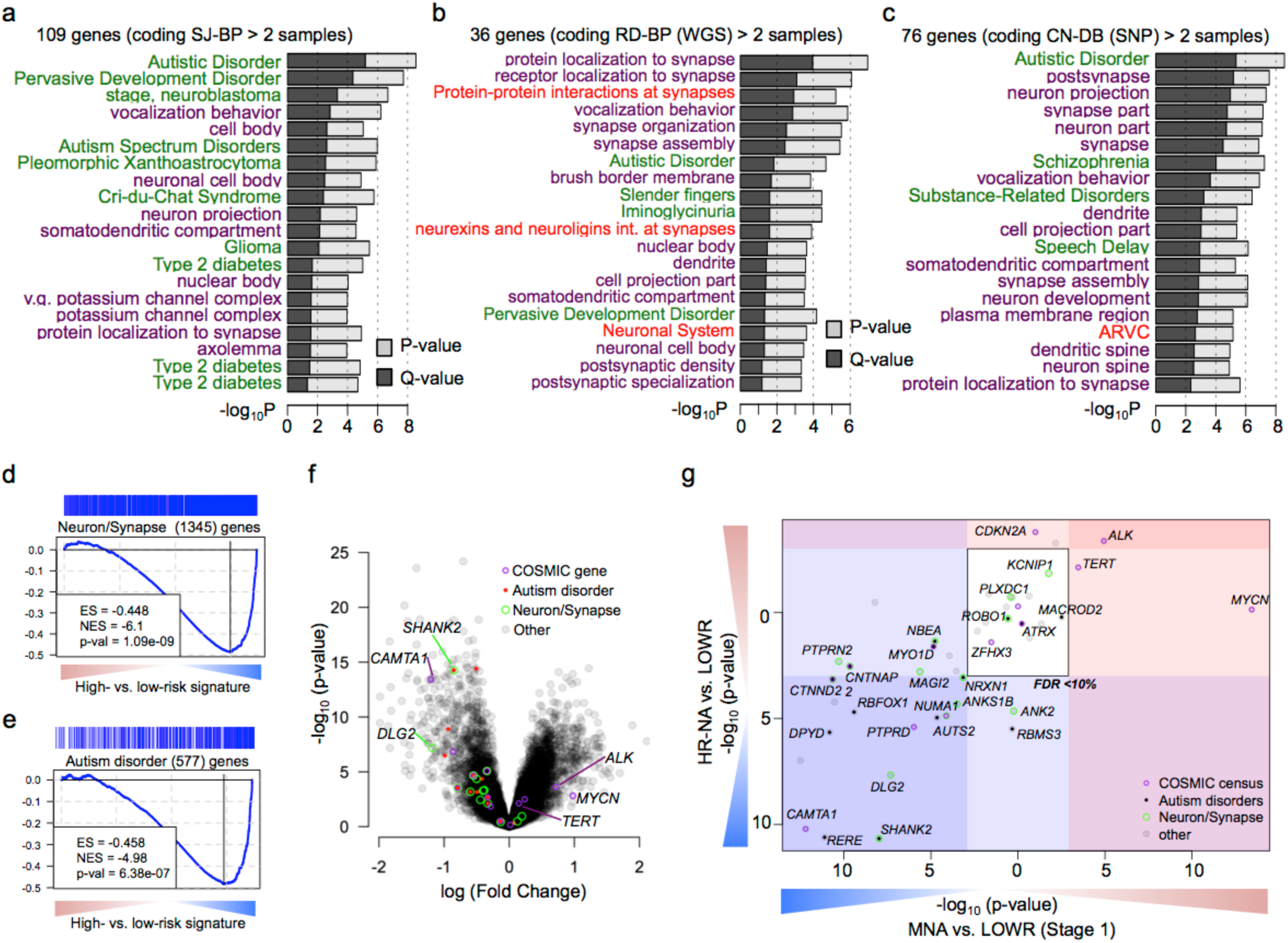
Neurodevelopmental genes are recurrently targeted by structural variations in neuroblastoma. (a-c) Function enrichment analysis bar plots for genes recurrently altered based on breakpoint analyses of (a) SJ-BPs, (b) RD-BPs and (c) CN-BPs. Analysis includes gene sets associated with diseases (green), Gene Ontology (purple) and Pathways (red). (d-e) Gene Set Enrichment Analysis across the signature of high-versus low-risk tumors from the HumanExon array show enrichment of (d) neuronal and synapse part and (e) autism disorder predisposition genes. (f) Volcano plot showing differential expression between high- and low-risk highlighting genes with recurrent SVs and their functional classification (g) Subtype specific high-versus low-risk differential expression analysis of 77 recurrently altered genes from Fig 4i shown as scatter plot (MNA = x-axis, HR-NA = y axis). (d-g) Analysis replicated in two datasets: HuEx arrays (here) and RNA-seq (Supplementary Fig. 19a-d).

### Recurrently disrupted neurodevelopmental genes are down-regulated in high-risk neuroblastoma

To further characterize the clinical relevance of recurrently altered genes in neuroblastoma, we studied their differential expression between high-risk subtypes and low-risk (Stage 1 and 4s) groups (**Fig. 4d,e**). We first used gene set enrichment analysis (GSEA)(Subramanian et al. 2005) to confirm the directionality of the regulation of gene classes enriched in recurrently altered genes. We observed down-regulation of neuronal and synaptic genes (P_HuEx_= 1.09 × 10^-9^) and autism disorder susceptibility genes (P_HuEx_= 6.38 × 10^-7^) in high-risk tumors when compared to stage 1 low-risk tumors (**Fig. 4d-f**). We then focused on differential expression of genes with recurrent SVs in high-risk subtypes (**Fig. 4g**). Known oncogenes including *TERT* and *ALK* were up-regulated in both MNA and HR-NA while *MYCN* was up-regulated only in MNA tumors. Known neuroblastoma tumor suppressor genes including *CAMTA1* and *RERE* from the 1p chromosome region and *PTPRD* were down-regulated in both subtypes. Most genes with a role in ASD predisposition, and those involved in neuron and synapse formation, were down-regulated in both high-risk subtypes. In particular, expression was significantly reduced for *SHANK2* (P_MNA_ = 2.15 × 10^-11^; P_HR-NA_= 1.05 × 10^-8^) and *DLG2* (P_MNA_ = 2.1 × 10^-8^;P_HR-NA_= 4.86 × 10^-8^) in high-risk compared with stage 1 low-risk tumors and compared to stage 4S low-risk tumors (P_MNA_ = 1.41 × 10^-3^;P_HR-NA_= 1.82 × 10^-5^ and P_MNA_ = 1.09 × 10^-4^; P_HR-NA_= 2.72 × 10^-4^ respectively). These results replicated using RNA-sequencing data (**Supplementary Fig. 19a-d**).

### Neurodevelopmental genes *SHANK2* and *DLG2* are frequently disrupted by chromosome 11 translocation events

High-risk neuroblastomas without *MYCN* amplification frequently exhibit deletion of chromosome 11q and this event is associated with a poor outcome (Guo et al. 1999; Attiyeh et al. 2005; Caren et al. 2010). The most frequent breakpoints observed in this study were located at chromosome 11q.13 and 11q.14 and disrupted the *SHANK2* and *DLG2* gene loci respectively (**Fig 5a,b; Supplementary Fig. 20**). *SHANK2* translocation partners involved chromosome 17q in 10/11 WGS cases, in addition we identified 49 samples from the SNP dataset (10.7%) with breakpoints in *SHANK2* (**Fig 5a**). *DLG2* translocation partners included multiple chromosomes. *DLG2* breakpoints were also identified in 28 samples from the SNP array dataset (**Fig 5b**).

**Figure 5:**
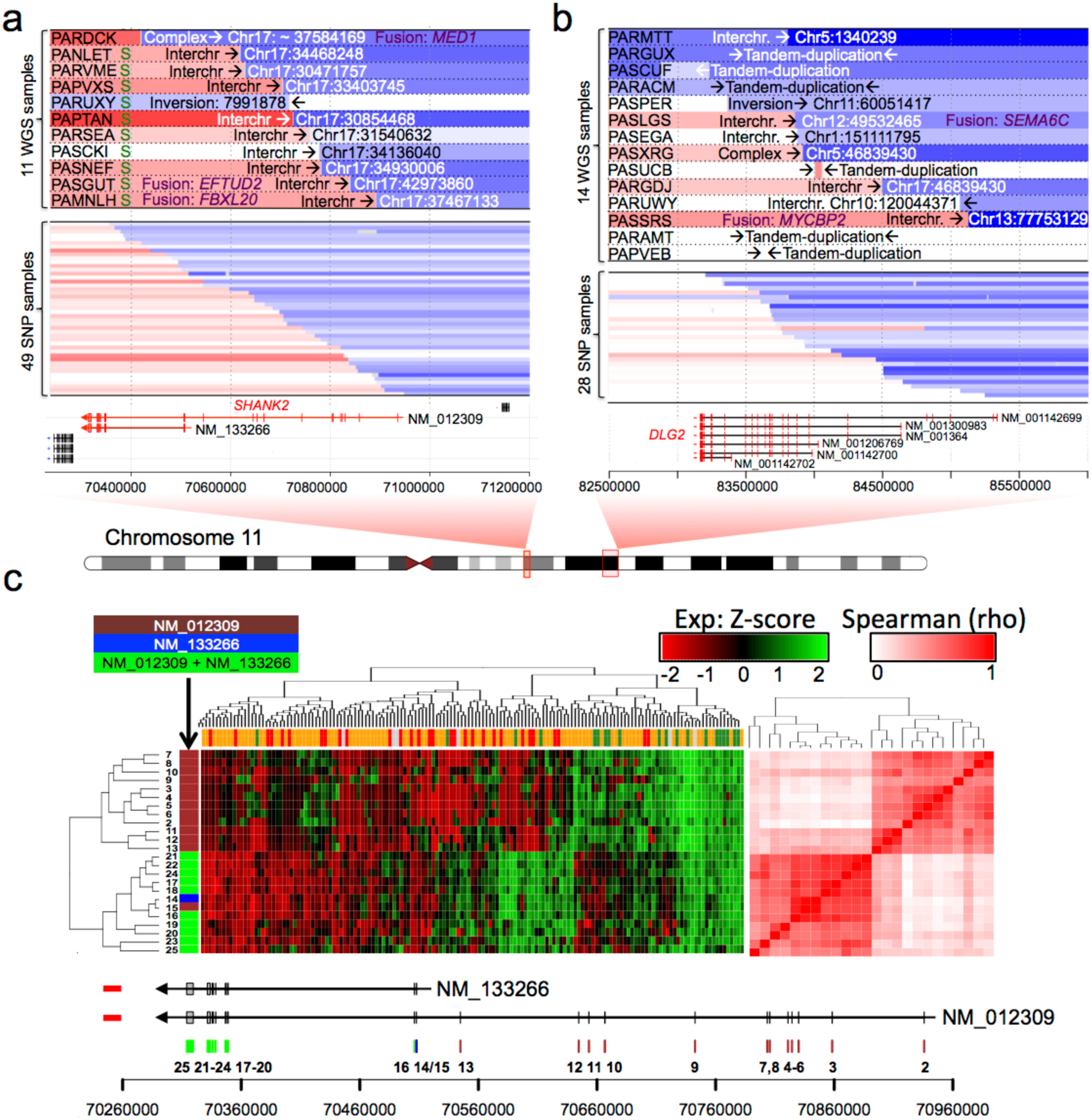
Neuronal genes *SHANK2* and *DLG2* are frequently disrupted by translocation events involving chromosome 11. **(a-b)** Copy number, breakpoint location and types of SVs at genomic regions harboring rearrangements that span (a) *SHANK2 and* (b) *DLG2* loci; ‘S’ at the left of the panel indicates positive validation by Sanger sequencing for *SHANK2* (Supplementary Fig. 15) and *DLG2* (Supplementary Fig 16). Associated gene fusion events obtained from RNA-seq indicated in purple text. **(c)** Clustering analysis of *SHANK2* exon level FPKM from RNA-seq data. The heatmap (left) shows higher exon expression level in S4s compared to MNA and HR-NA samples. The correlation matrix (right) shows two well-defined clusters associated with the two known coding isoforms of the gene. Exons are color coded according to their isoform span.

*SHANK2* is a scaffold protein in the postsynaptic density (PSD) with two known coding isoforms (long: NM_012309; short: NM_133266). We therefore studied the expression pattern of *SHANK2* at the exon level using both RNA-seq (**Fig. 5c**) and HumanExon arrays (**Supplementary Fig. 21a,b**) data. Clustering analysis of *SHANK2* exon expression revealed two distinct clusters corresponding to the two known coding isoforms. Expression of the long isoform (NM_012309) was decreased in high-risk tumors compared to INTR and LOWR as observed from RNA-seq (**Fig. 5c**) and HuEx expression analysis (**Supplementary Fig. 21a,b**). Finally, in a large independent RNA-Seq cohort (Wang et al. 2014a), reduced expression of the long isoform (NM_012309) was associated with increased tumor stage (P=1.62 × 10^-22^, **Supplementary Fig. 21c**) and poor overall survival (P=7.21 × 10^-13^, **Supplementary Fig. 21d**). Notably, the association with poor survival remained significant even within the low- and intermediate-risk subsets of neuroblastoma that typically have favorable outcomes (P=2.22 × 10^-5^, **Supplementary Fig. 21e)**. Consistent with the *SHANK2* expression pattern, we observed decreased activation of PSD genes based on GSEA in high-risk compared to low-risk neuroblastomas in multiple prognostic signatures (**Supplementary Fig. 22**). We decided to further study the long isoform of *SHANK2* (NM_012309) given that nearly all SVs uniquely disrupt this splice variant, leaving the short isoform (NM_133266) intact.

### *SHANK2* expression inhibits cell growth and viability of neuroblastoma cells

To further elucidate the role of *SHANK2* in neuroblastoma, three neuroblastoma cell lines with low or no endogenous *SHANK2* expression (**Supplementary Fig. 23**) were selected. These included SY5Y (*MYCN* Non-amplified), Be(2)C (*MYCN* amplified), and NGP (*MYCN* amplified). Cells were stably transduced to constitutively overexpress *SHANK2* long isoform or an empty vector control. SHANK2 expression was confirmed by Western blot (**Fig. 6a-c**). When maintained in selection media and grown alongside empty vector controls, the SHANK2-expressing cells consistently exhibited decreased cell growth and viability as measured by RT-CES cell index (**Fig. 6d-f**) as well as CellTiter Glo assay (**Fig. 6g-i**). For SY5Y, when control reached confluence, the comparable cell indexes of the SHANK2 overexpressing lines were reduced by 75% (P=3.4 × 10^-5^; **Fig. 6d**), Be(2)C cell index reduced by 62% (P=3.16 × 10^-4^; **Fig. 6e**), and NGP showed a 14% reduction (P=2.62 × 10^-2^; **Fig. 6f**). We also observed decreased cell viability in SHANK2-expressing cells at both 4- and 7-day endpoints using an ATP-dependent CellTiter Glo assay. Specifically, viability of SY5Y SHANK2-expressing cells was reduced to 65.51% (P=1.34 × 10^-18^) and 52.64% (P=4.72 × 10^-26^) of controls (**Fig. 6g**). This was reinforced in the similar results for Be(2)C SHANK2-expressing cells (49.21% and 44.26%, P=5.76 × 10-^28^ and 5.74 × 10^-15^; **Fig. 6h**) and NGP (90.63% and 74.01%, P=5.11 × 10^-3^ and 6.01 × 10^-13^) (**Fig. 6i)**.

**Figure 6:**
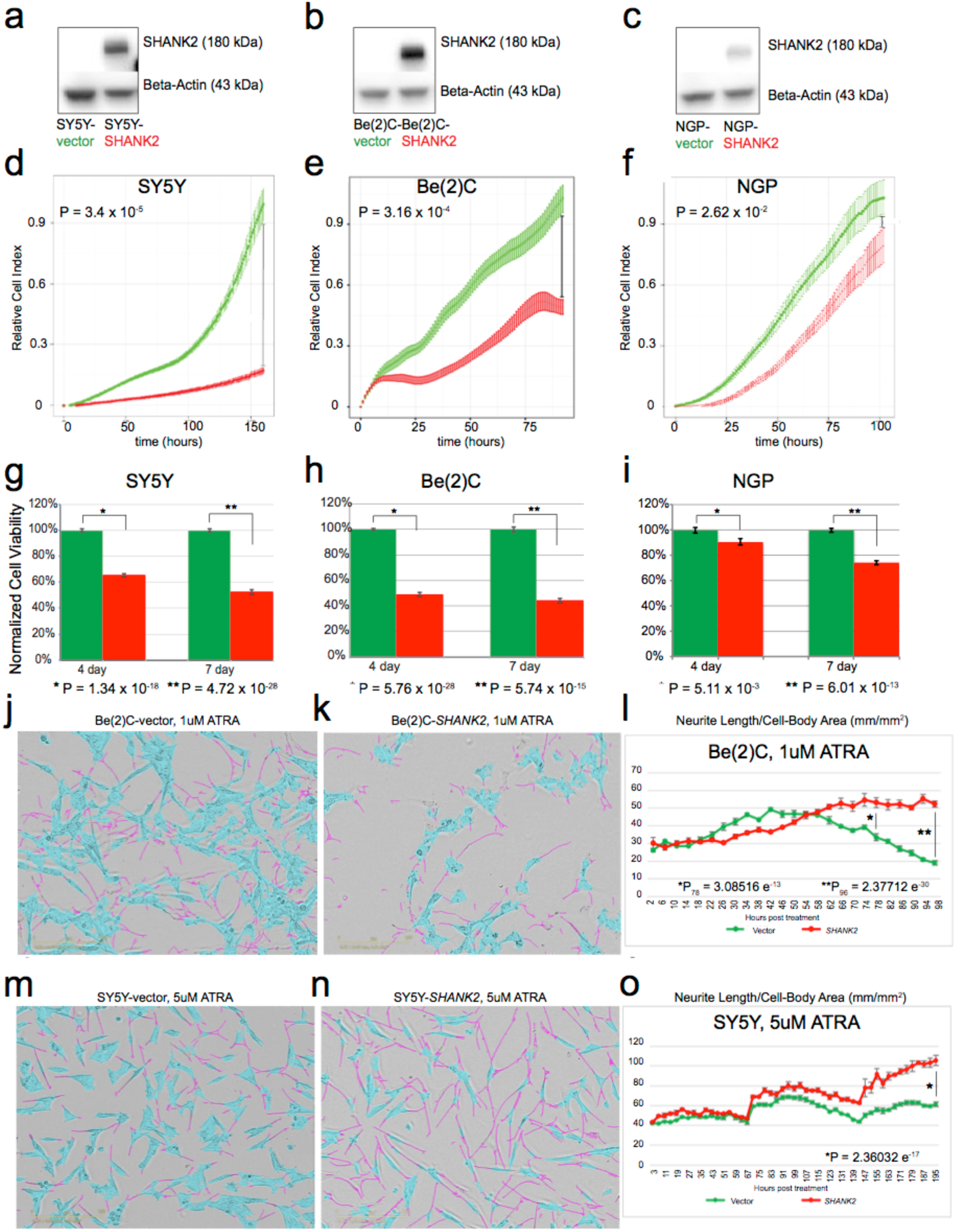
*SHANK2* reduces cell growth and promotes differentiation in neuroblastoma cell line models. (a-c) Western blots confirming overexpression of SHANK2 in all tested neuroblastoma cells: (a) SY5Y, (b) Be(2)C and (c) NGP. (d-f) Decreased proliferation in all 3 lines overexpressing SHANK2 (red) compared to controls (green), as measured by RT-CES. (h-i) Decreased viability in SHANK2 overexpressing cells (red) versus controls (green) as measured by ATP-dependent CellTiter Glo Assay. (j,k) Incucyte images Be(2)C cells for (j) vector control and (k) SHANK2 expressing cells at 78 hours post treatment with 1 uM ATRA. Neurite extensions masked in pink; cell bodies masked in blue. (l) Neurite length normalized to cell body area starting immediately after ATRA application corresponding to Be(2)C cells images at different time points. (m, n) SY5Y images from Incucyte at day 9 post ATRA treatment (5uM). (o) Neurite outgrowth normalized to cell body area in corresponding to SY5Y cells images at different time points.

### *SHANK2* expression accelerates differentiation of neuroblastoma cells exposed to all-trans retinoic acid (ATRA)

We next investigated the role of *SHANK2* in neuronal differentiation in Be(2)C and SY5Y cells exposed to ATRA. In the presence of ATRA, overexpression of *SHANK2* accelerated differentiation as measured by presence and length of neurites compared to cell body (**Fig 6j-o**; **Supplementary Fig. 24a-f**). While decreases in growth can be measured even without drug application, once ATRA is applied, cells overexpressing *SHANK2* developed neurites more quickly, and those neurites extended further than observed in empty vector controls (**Supplementary Fig. 24c,d**). In Be(2)C cells, a significant difference in neurite outgrowth normalized to cell-body area was seen at 72 hours post treatment with 1 uM ATRA (**Fig. 6j-l**), with *SHANK2* cells exhibiting a 1.6-fold increase over controls (P=3.09 × 10^-13^); the difference increased to 2.76-fold at 96 hours (P=2.37 × 10^-30^). Even with vehicle alone, *SHANK2* cells had more neurite outgrowth per cell body compared to their empty vector counterparts at both 72 and 96 hours post treatment (P=1.02 × 10^-5^, P=1.25 × 10^-13^, respectively). In SY5Y, though differentiation took longer and both *SHANK2* cells and controls eventually reach 100% confluence with vehicle alone, SHANK2 overexpression still decreased confluence (P=1.69 × 10^-6^, **Supplementary Fig. 24b**). In analyzing total neurite outgrowth without normalization for cell body area, SY5Y ATRA-treated *SHANK2* cells outpaced controls st*a*rting at hour 144 post-treatment and continued to lead until the experiment ended, with a total neurite measurement increased 1.55-fold over controls (P=1.62 × 10^-35^; **Supplementary Fig. 24d**). Once normalized, *SHANK2* cells have higher measured outgrowth starting at 75 hours post treatment through hour 96, and maintain from there. At 195 hours past treatment, *SHANK2* cells treated with 5uM ATRA displayed neurites at 1.71-fold increase over their empty vector controls (P=2.36 × 10^-17;^ **Fig. 6m-o**). Taken together, these data suggest *SHANK2* is a newly identified haplo-insufficient tumor suppressor in high-risk neuroblastoma that is disrupted by recurrent somatic structural variation in the *MYCN* non-amplified subset of cases.

## DISCUSSION

Sequencing studies of neuroblastoma tumors have revealed a relatively low SNV burden and limited mutational landscape (Cheung et al. 2012; Molenaar et al. 2012; Pugh et al. 2013; Sausen et al. 2013), leaving aneuploidy and large segmental chromosomal alterations as the main candidate driver mutations in many tumors. Structural variations (insertions, deletions, duplications and translocations, and inversions) can also function as potent cancer drivers, as demonstrated with the discovery of rearrangements near the *TERT* gene driving aberrant telomerase expression in many high-risk neuroblastomas (Peifer et al. 2015; Valentijn et al. 2015). Here, we expand the landscape and understanding of structural variation in neuroblastoma through an integrative genomic analysis of a large cohort of patient samples profiled by whole genome sequencing and SNP arrays together with additional transcriptional data. To the best of our knowledge, this study represents the largest integrated genome-wide survey of structural variation in neuroblastoma including alignment based (SJ-BP) and copy number based (RD-BP and CN-BP) structural variation breakpoint analyses.

Structural variation complexity was most evident in high-risk tumors without amplification of *MYCN* (HR-NA). This observation is consistent with other reports of increased chromosomal instability in this high-risk subset (Caren et al. 2010). The current study strengthens and extends previous reports of chromosomal instability by including additional structural variation types and breakpoint burden analyses. Specifically, we observed significantly more tandem-duplications HR-NA tumors. These affected nearly every chromosome, except for chromosome 2, which contains the *MCYN* amplicon. HR-NA tumors harbored more SVs in known cancer genes as well as novel genes. In contrast, the SNV burden was very similar between MNA and HR-NA groups. As suggested by pan-cancer studies, the underlying mechanisms potentiating chromosomal instability and somatic SNV burden may differ (Ciriello et al. 2013). The mechanism leading to the observed increase in chromosomal instability in HR-NA tumors remains unknown. Pan-cancer studies have reported *TP53* mutations as a major driver of chromosomal instability; however, *TP53* loss of function is rarely observed in neuroblastoma at diagnosis.

Chromothripsis has been reported in as many as 18% of high-stage neuroblastomas (Molenaar et al. 2012). Similarly, in the current study, 19% of high-risk tumors from the TARGET cohort exhibit chromothripsis (N=20/105), involving a total of 27 chromosomal regions. These events largely overlap with amplification of *MYCN* (as well as some *ALK* cases) on chromosome 2p and *TERT* on chromosome 5p, suggesting an important role of chromothripsis followed by purifying selection as an underlying cause of those alterations. We also observed high-breakpoint density in the X chromosome of females based on the SNP array data, which could be explained by higher tolerance to chromothripsis in diploid regions. Future studies are required to determine whether the oncogenic role of chromothripsis represents an opportunity for therapeutic intervention.

Along this study, we report a shared repertoire of genes altered in neuroblastoma and neurodevelopmental disorders (NDD), including autism spectrum disorder. A link between cancer and autism has been previously established in PTEN-associated germline syndromes (Goffin et al. 2001), and multiple autism susceptibility genes also have a known role in cancer (Crawley et al. 2016). Moreover, certain germline deletions associated with NDD, such as 10p15 (DeScipio et al. 2012) and 16p24.3 (Willemsen et al. 2010), are reported here to occur somatically in neuroblastoma. Transcriptomic analyses have shown that neural lineage pathways are commonly down-regulated in high-risk neuroblastomas compared to low-risk signatures (Fredlund et al. 2008). Here, we show that structural variation preferentially disrupts neurodevelopmental genes in neuroblastoma. We hypothesize that structural variations in *SHANK2* and other coding proteins of the postsynaptic density (PSD) comprise novel neuroblastoma candidate tumor suppressors involved in neuronal differentiation. Additional candidate genes identified here with a proposed role in neurotransmission and synapsis and involvement in autism include *DLG2, AUTS2, CNTNAP2, NRXN1, CTNND2(Gai et al. 2012)*. Structural variants affecting these genes are more prevalent in high-risk neuroblastomas without amplification of *MYCN*, which is itself a potent driver of dedifferentiation(Westermark et al. 2011).

We propose that disruption and deregulation of *SHANK2* promotes the undifferentiated state of neuroblastoma cells, and that other synaptic genes may play a similar role in this childhood cancer. Synaptogenesis is a key process in neuronal differentiation and mutations in the *SHANK* family of proteins are frequently implicated in NDD (also termed shankopathies), offering potential therapeutic opportunities for these disorders (Wang et al. 2014b). In the current study, we show that *SHANK2* is disrupted by recurrent somatic SVs in HR-NA tumors and that *SHANK2* expression is low across high-risk tumors. The mechanism driving low *SHANK2* levels in the MNA high-risk tumors remains to be identified. We further demonstrate that decreased *SHANK2* expression is associated with poor survival in neuroblastoma, even within the low- and intermediate-risk subsets of patients that typically have good outcomes. This suggests that *SHANK2* expression, or the undifferentiated cell state, may serve as a biomarker for non-high-risk tumors requiring more aggressive treatment. Our *in vitro* studies demonstrate that forced expression of *SHANK2* reduces cell growth and increases neurite outgrowth (indicative of differentiation) in human derived neuroblastoma cell lines exposed to ATRA. Given that retinoids are currently utilized as maintenance therapy in high-risk neuroblastoma standard of care (Matthay et al. 1999; Matthay et al. 2009), the sensitizing effect of *SHANK2* expression to ATRA treatment underscores the importance of understanding the mechanisms driving and maintaining the undifferentiated phenotype of high-risk neuroblastoma. Subsequent studies with larger cohorts should evaluate the role of mutated and deregulated neurodevelopmental genes in retinoic acid treatment response. Taken together, we depict a substantially expanded landscape of structural variation in neuroblastoma and provide mechanistic insight into the aberrant neuronal development hallmark of the high-risk form of this childhood cancer.

## METHODS AND DATA ACCESS

### Datasets and data availability

The primary dataset in this study is comprised of 135 whole genome sequenced tumor-blood matched pairs (Complete Genomics (CGI)) and 914 SNP-array profiled tumor-blood pairs (Illumina SNP arrays). CGI short read sequencing uses commercial software for processing, aligning to reference genome (hg19) and variant calling (Cancer Pipeline 2.0; http://www.completegenomics.com/documents/DataFileFormats_Cancer_Pipeline_2.0.pdf). Matched RNA sequencing (n=153) and Affymetrix HumanExon array data (n=247) were analyzed to assess the influence of SVs on mRNA expression. Whole genome and RNA-sequencing data are available from dbGaP (https://www.ncbi.nlm.nih.gov/gap) with study-id phs000218 and accession number phs000467. The SNP array data generated for this study has been uploaded to Gene Expression Omnibus (GSE131189). Processed data from CGI whole genome sequencing (including SV and SNV calls) and the HumanExon arrays are available from the TARGET data NCI data matrix (https://target-data.nci.nih.gov/). The SEQC neuroblastoma RNA-seq dataset used for survival analyses is available from Gene Expression Omnibus (GSE62564). Additional genomic profiles from neuroblastoma cell lines are available as follows: SNP genotyping arrays (Gene Expression Omnibus: GSE89968) and RNA-seq (Gene Expression Omnibus: GSE89413)(Harenza et al. 2017).

### Copy number segmentation, visualization (IGV) and recurrence analysis

CGI “somaticCnvDetailsDiploidBeta” (https://target-data.nci.nih.gov/Controlled/NBL/WGS/CGI/) files provide information on estimated ploidy and tumor/blood coverage ratio for every 2-kb along the genome. We used custom scripts to reformat coverage data to be processed with “copynumber” R bioconductor pakage(Nilsen et al. 2012). We then utilized Winsorization (winsorize) data smoothing and segmentation with piecewise constant segmentation (pcf) algorithm with attributes kmin=2 and gamma=1000. Segmented data was visualized with IGV and further used as input to GISTIC2.0(Mermel et al. 2011). GISTIC attributes were as follows: -v 30 -refgene hg19 -genegistic 1 - smallmem 1 -broad 1 -twoside 1 -brlen 0.98 -conf 0.90 -armpeel 1 -savegene 1 -gcm extreme -js 2 -rx 0.

### Filtering of CGI SV calls

The CGI Cancer Pipeline 2.0 produces a full report including quality control, variant calling and CNV analyses. The “somaticAllJunctionsBeta” files provide information for individual junctions detected in a tumor genome that were absent in the corresponding normal genome. The “highConfidenceJunctionsBeta” files contain a filtered subset of the junctions reported in “somaticAllJunctionsBeta” file. This subset includes junctions that likely resulted from a true physical connection between the left and right sections of the junctions. A detailed description of the approach followed by the vendor (CGI) is available in the Data File Formats description (http://www.completegenomics.com/documents/DataFileFormats_Cancer_Pipeline_2.0.pdf). Additional filtering of high confidence variants was initially applied to the entire TARGET repertoire of tumor datasets including ALL, AML, NBL, OS, CCSK and RT in order to remove recurrent junctions consistent with common variation. We added additional filters to remove rare/common germline variants that passed CGI filters as well as artifacts and low confidence variants. To this end, we used the Database of Genomic Variants (DGV v. 2016-05-15, GRCh37) in order to remove SVs which reciprocal overlap with DGV annotated common events was higher than 50%. We only filtered variants which type matched in both CGI SV set and DGV database.

### Mapping and annotation of filtered alignment-based SVs (SJ-BP)

We used RefSeq gene definitions for hg19 downloaded from UCSC (10/31/2018) in order to map SV calls to nearby genes. First, we used two approaches to map variants: numerical changes (tandem-duplications and deletions size <2Mb) containing whole genes were used to define copy number alterations. Second, we mapped breakpoints relative to gene exonic coordinates. SVs were considered ‘disrupting’ when either one of the breakpoints localized between transcription start and ends of any of isoforms of a gene and ‘proximal’ when localized within 100Kb upstream or 25Kb downstream the most distal isoform of each gene; a graphical description is represented in **Figure 4b**. In addition, we considered ‘intronic’ SVs as those in which both breakpoints mapped to the same intron.

### Processing DNA copy number segmentation from WGS

We processed ‘cnvDetailsDiploidBeta’ files from the Cancer Pipeline 2.0 (CGI^©^) containing average normalized read-depth coverage values at every 2-Kb sliding window throughout the genome. Then, tumor/blood normalized ratios were subject to piecewise constant segmentation algorithm(Nilsen et al. 2012) implemented in the ‘copynumber’ R package. The processed segmentation file is available through the TARGET data matrix (https://target-data.nci.nih.gov/).

### Generating DNA copy number segmentation from SNP arrays

We genotyped 914 matched patient tumor and normal samples using Illumina SNP arrays. This cohort included 488 samples which were previously reported(Attiyeh et al. 2009) and reanalyzed here. The complete dataset comprised three different genotyping architectures from Illumina: HumanHap550, Human610-Quad, and HumanOmniExpress. The common set of 316,210 SNPs were processed for tumor DNA segmentation using the *SNPrank* algorithm implemented by the NEXUS^®^ software platform. The generated segmentation data for is 914 unique samples is available through the TARGET data matrix (https://target-data.nci.nih.gov/).

### Mapping and annotation of DNA amplifications, deep deletions and copy number breakpoints from WGS (RD-BP) and SNP arrays (CN-BP)

Breakpoints were called from segmentation profiles. Due to artifacts observed at subtelomeric and pericentromeric regions, these regions were excluded. For the rest of the genome, a breakpoint was called when the absolute value of the copy number log-ratio difference between contiguous segments is higher than 0.152 for SNP arrays and 0.304 for WGS; both cutoffs account for 10% and 20% copy number change respectively (i.e for diploid regions, ΔCN = 0.2 and ΔCN = 0.4 respectively). For the WGS dataset, we called amplifications when CN >=8 and deep deletions when CN <= 0.5. For SNP arrays, we used less stringent cutoffs for amplification (CN >= 4.5) and deep deletions (CN <= 0.9). These relaxed criteria were selected due to SNP arrays having a narrower dynamic range and lower resolution. We used RefSeq gene definitions for hg19 downloaded from UCSC (October 31^st^, 2018 version) to map copy number alterations and breakpoints to nearby genes. Genes were considered amplified or deep deleted when all isoforms were contained within the altered segment boundaries. Breakpoints were considered ‘disrupting’ when the breakpoint localized between transcription start and ends of any isoform of a gene and proximal when localized at within 100Kb upstream or 25Kb.

### Co-localization of breakpoints derived from WGS (SJ-BP & RD-BP) and SNP arrays (CN-BP)

We studied the co-localization of breakpoints derived from alternative measurements. First, we compared SJ-BPs and RD-BPs in each of the 135 WGS samples; overall, 30.5% of SJ-BPs co-localize with a RD-BPs (**Supplementary Fig**.**3a**) whereas 62% RD-BPs matched with SJ-BPs (**Supplementary Fig. 3b**). The lower overlap in SJ-BPs is expected since not all SVs necessarily involve a change in copy number dosage (i.e. inversions and translocations). We next evaluated the co-localization of breakpoints across WGS and SNP platforms within the subset of 52 overlapping samples. 50.2% of CN-BPs from SNP arrays co-localized with SJ-BPs from the WGS dataset (**Supplementary Fig**.**3c**) whereas only 8.2% of SJ-BPs co-localize with CN-BPs (**Supplementary Fig. 3d**). Furthermore, when comparing dosage based breakpoints across platforms (RD-BP and CN-BP), 23.6% RD-BPs where found co-localizing CN-BP (**Supplementary Fig. 3e**) whereas 66.6% CN-BP co-localized with RD-BPs (**Supplementary Fig. 3f**). Overall, SNP arrays display reduced the number of breakpoints compared read-depth based profiles; we attribute these differences to a narrower dynamic range and lower probe density of the platform. Finally, we performed a randomized test by sample shuffling (N_i_=1000) in order to evaluate whether each of the co-localization percentages listed above could arise by chance or due to recurrence of SVs across samples. All randomized percentage distributions range between 0.7% and 2.3%; in all cases the null hypothesis was discarded (p-value < 0.001, **Supplementary Fig. 3g-l**). Taken together, alternative breakpoint detection methods returned consistent results even when derived from different platforms providing means for both orthogonal and cross-platform validation of SVs. However, certain types of SVs can only be detected using alignment-based methods.

### Tumor mutational burden analyses

We obtained measures representative of the burden of different mutation types under study (including SNVs, SVs and BPs). To this end, the density of mutations of every type is calculated as the average number of mutations in a given sample per sequence window (10Mb for SVs and BPs and 1Mb for SNVs). Instead of a single density value per sample we measure mutational densities for each chromosomal arm, excluding short arms with very low mappability (13p, 14p, 15p, 21p, 22p and Y chromosome). The remaining 41 chromosomal arms in each sample represent single sample distributions of mutational densities from which quantiles are obtained. We used the interquartile mean (IQM) since it offered a measure robust against outliers while conserving the variability across samples even in low-density breakpoint samples.

### Filtering and annotation of likely pathogenic somatic SNVs

The CGI cancer pipeline 2.0 provides somatic variant calls for SNVs and small indels. Given the gapped nature of CGI reads which leads to high noise to signal ratio, we incorporated additional SNV filtering. We first annotated CGI SNV calls using Variant Effect Predictor (VEP) pipeline(McLaren et al. 2016). Our filter follows two steps: 1) collect high quality somatic non-synonymous coding variants (Phred like Fisher’s exact test P<0.001) annotated as having a moderate or high functional impact; this set of variants was combine with COSMIC catalogue of pathogenic variants (release v84). 2) Hot-spot analysis of variants from our combined catalogue (step 1) to identify both clonal and low allele frequency pathogenic variants.

### Gene fusion analysis

Gene fusion analysis from RNA-seq data was studied using three available tools: (STAR-fusion(Dobin et al. 2013), fusionCATCHER(Daniel Nicorici 2014) and DeFUSE (McPherson et al. 2011)). We then collected fusion events that matched inter-chromosomal events from the CGI structural variation calls.

### Statistical and survival analyses

All statistics performed with genomic data Wilcoxon rank-sum test, Kruskal-Wallis test and survival analyses were done using R programming. For the survival analyses, we used the R ‘survival’ library. In order to estimate the association between *SHANK* long isoform expression and survival we obtained the optimal separation (lower log-rank test p-value) from all possible expression thresholds and then used Benjamini & Hochberg (false discovery rate) for multiple testing correction and q-value estimation.

### Sanger sequence validation

From alignment and variant calls provided by Complete Genomics, SV breakpoints were mapped and junction sequence was assembled using public UCSC Genome browser. The assembled sequence was then submitted into Primer3 to engineer PCR primers to bridge the breakpoints. Resultant primers were then checked against BLAT as well as an internal algorithm for binding specificity. PCR reactions were then carried out on 25 ng of DNA using optimized conditions for each reaction. Products were checked via gel electrophoresis for specificity to expected size and uniqueness. If the product had multiple bands, the entire remaining sample would be run out then bands of interest excised and the DNA extracted using MinElute Gel Extraction Kit from Qiagen. Products with single bands were cleaned up and prepared for sequencing using the MinElute PCR Purification Kit (Qiagen). Samples were then sequenced at a core facility with 2 picomoles of the same primer used to create amplicon. Resultant sequences were then aligned to the expected sequence assembled from CGI results and mapped to chromosomes using BLAT at UCSC genome browser.

### Cell culture

Cells were grown in RPMI-1640 with HEPES, L-glutamine, and phenol red (cat # 22400-089), supplemented with 10% Fetal Bovine Serum, 1% antibiotic-antimycotic (cat # 15240-062), and 1% L-glutamine (cat # 25-005-CI) in 5% CO_2_ at 37°C in the dark. Transduced cells also had the appropriate concentration of puromycin in media for selection.

### LentiVirus infection

Lentiviral vector plasmid for the long isoform of *SHANK2* (NM_012309) was obtained commercially from GeneCopoeia (EX-H5274-Lv105). Empty vector control plasmid pLv105 was originally from GeneCopoeia. Creation of the virus media was accomplished using Lipofectamine 3000 ™ applied to 293TN cells with packaging plasmid psPAX2, envelope plasmid pMD2.g, and the Lentiviral backbone plasmid containing the ORF for NM_012309 or empty vector. Infectious viral media was pooled over 2 days then filtered through 0.45µm nitrocellulose and combined with polybrene at 8 µg/mL media and applied to cells. Following infection, transduced cells were selected with puromycin in line-dependent concentrations.

### Growth and proliferation assay using RT-CES

Cells were plated in 96-well RTCES microelectronic sensor arrays (ACEA Biosciences, San Deigo, CA, USA). Density measurements were made every hour. Cell densities were normalized to 5 hours post-plating.

### Cell viability assays

Cells were plated in clear-bottomed, 96-well plates in 200 µL media and allowed to grow under normal conditions for either 4 or 7 days. Before reading, 100 µL media was replaced with equal volume of CellTiter Glo ® reagent and read on a GloMax Multi-detection instrument (Promega). Arbitrary luminescence units were normalized to empty vector-transduced controls and results expressed as percentages of control levels from the same assay.

### ATRA-induced differentiation

Cells were plated in normal media at optimized densities for each parental line in 96-well plates and allowed 24-48 hours to firmly attach to plates. Media was then switched for low-serum media containing either 1% or 3% FBS and allowed 24 hours to equilibrate, after which it was replaced with low-serum media supplemented with varying concentrations of ATRA (all-*trans*-retinoic-acid, Sigma, R2625) or vehicle (DMSO) alone, in volume corresponding to the highest concentration of ATRA for each experiment. Plates were then left in normal growth conditions and protected from light. RA media was refreshed every 72 hours to prevent oxidation. Plates were placed in an IncuCyte ZOOM™ instrument to utilize live cell imaging. Each well was imaged every four hours and the “NeuroTrack” software module to quantify neurite outgrowth.

### Protein isolation and Western blotting

Whole cell lysates were created by applying denaturing lysis buffer containing protease/phosphatase inhibitors (Cell Signaling Technology, 5872) to cells on ice and allowing lysis for 30 minutes. The total sample was sonicated for 5 seconds and spun at max speed in a microcentrifuge for 15 minutes at 4°C before collecting supernatant to clean tube. Quantification of protein was done using the Pierce BCA Protein Assay Kit (Thermo, 23227). Protein was loaded on 4-12% Tris-Glycine gels, transferred to PVDF membrane, and probed with antibodies in 5% milk in TBST. Antibody stripping used Restore ™ Stripping buffer (Thermo, 21059). Detection of HRP-conjugated secondary antibodies used SuperSignal™ West Femto Maximum Sensitivity Substrate (Thermo, 34096).

## AKNOWLEDGEMENTS

This work was supported in part by NIH grant R01-CA124709 (S.J.D) and the Roberts Collaborative Forefront Award (G.L.). This project was also funded in part by a supplement to the Children’s Oncology Group Chair’s grant CA098543 and with federal funds from the National Cancer Institute, National Institutes of Health, under Contract No. HHSN261200800001E to S.J.D and Complete Genomics.

## AUTHOR CONTRIBUTIONS

S.J.D designed the experiment. G.L. and S.J.D. drafted the manuscript. G.L. and S.J.D. performed analyses of SVs from WGS. G.L. performed RNA data analysis. G.L. and A.M. performed *de novo* transcript analyses. G.L. and K.S.R. performed fusion transcript analyses. K.L.C. and M.D. performed Sanger sequencing. K.L.C., M.D., L.M.F., and E.H. performed *SHANK2* experiments. Z.V. assisted with sequence data analysis. J.S.W. and J.K. generated RNA sequencing data. S.A. and R.C.S. generated array-based expression data. All authors commented on or contributed to the current manuscript.

## ONLINE SUPPLEMENTAL MATERIAL

Supplementary data include 10 tables and 24 figures.

