## Supplementary Figures for "Somatic structural variation targets neurodevelopmental genes and identifies *SHANK2* as a tumor suppressor in neuroblastoma"

### I. SUPPLEMENTARY TABLES

**Supplementary Table 1: Neuroblastoma patient clinical and biological characteristics.**

| Covariate | WGS (CGI) | SNP (Illumina) | RNA-seq (Illumina) | Expression array (Affy-HuEx) |
|---|---|---|---|---|
| **INSS Stage** | **105** (Stage 4)<br>**6** (Stage 3)<br>**1** (Stage 2B)<br>**23** (Stage 4S) | **650** (Stage 4)<br>**98** (Stage 3)<br>**26** (Stage 2B)<br>**16** (Stage 2A)<br>**82** (Stage 1)<br>**37** (Stage 4S)<br>**5** (Unknown) | **126** (Stage 4)<br>**6** (Stage 3)<br>**21** (Stage 4S) | **214**(Stage 4)<br>**1**(Stage 3)<br>**30** (Stage 1)<br>2 (Unknown) |
| **MYCN amplification status** | **29** (Amp)<br>**106**(Not-Amp)<br>**1**(Unknown) | **241** (Amp)<br>**670**(Not-Amp)<br>**3**(Unknown) | **31** (Amp)<br>**121** (Not-Amp)<br>**1**(Unknown | **60** (Amp)<br>**185** (Not-Amp)<br>**2** (Unknown) |
| **COG Risk Group** | **106** (High-risk)<br>**14** (Inter-Risk)<br>**15** (Low-risk) | **695** (High-risk)<br>**70** (Inter-Risk)<br>**145** (Low-risk)<br>**4** (Unknown) | **127** (High-risk)<br>**13** (Inter-Risk)<br>**13** (Low-risk) | **215** (High-risk)<br>**30** (Low-risk)<br>**2** (Unknown) |
| **Vital status** | **74** (Alive)<br>**61** (Dead) | **454** (Alive)<br>**353** (Dead)<br>**107** (Unknown) | **75** (Alive)<br>**78** (Dead) | **105** (Alive)<br>**140**(Dead)<br>**2**(Unknown) |
| **Age at diagnosis group** | **32** (< 18 mo/old)<br>**103** (>18 mo/old) | **265** (< 18 mo/old)<br>**646** (>18 mo/old)<br>**3** (Unknown) | **29** (< 18 mo/old)<br>**124** (>18 mo/old) | **32** (< 18 mo/old)<br>**215** (>18 mo/old) |
| **Gender** | **52** (Female)<br>**83** (Male) | **408** (Female)<br>**506** (Male) | **64** (Female)<br>**89** (Male) | **106** (Female)<br>**141** (Male) |

**Supplementary Table 2: Cohort clinical data and sample description (xls file)**
A survey of all samples used in this study including clinical information and availability for each genomic profiling platform.

**Supplementary Table 3: Filtered CGI structural variants calls (xls file)**
Survey of genes affected by structural variants (SVs). SVs are classified into disrupting and proximal. Gene list divided in three groups: Sheet 1, driver gene list contains high frequently and COSMIC cancer census altered genes. Sheet 2: low frequency (<4 samples) altered genes. Sheet 3. Manually filtered artifacts. See Online Methods.

**Supplementary Table 4: Chromothripsis and high-breakpoint density chromosomes (xls file)**
Quantification of different measures of structural variations per chromosomal arm used to infer chromothripsis in WGS samples.

**Supplementary Table 5: Recurrently altered genes by alternative breakpoint analyses (xls file)**
Survey of genes affected by structural variants (SVs) obtained by alternative breakpoint analyses and cohorts (SJ-BP, RD-BP and CN-BP). Altered genes are classified into different lists according to SV impact on “coding” or “non-coding” regions at each gene loci.

**Supplementary Table 6: Orthogonally identified recurrently altered genes (xls file)**
List of 77 genes disrupted by structural variants (SVs) with alignment (SJ-BP) and read-depth (RD-BP) based evidences from the WGS dataset.

**Supplementary Table 7: SNVs mapping to recurrently altered genes (xls file)**
Pathogenic non-synonymous SNVs found in the coding regions and splice sites of recurrently altered genes. Mutation Annotation Format (MAF) data file.

**Supplementary Table 8: Primer sequences used for Sanger sequencing validations (xls file)**
The table informs about sequences used for Sanger validation of SVs of *ALK*, *ATRX*, *DLG2*, *SHANK2* and *TERT*; includes Junction Id which relates to variants described in Supplementary Table 3.

**Supplementary Table 9: Combined identification of gene fusion transcripts and structural variations (xls file)**
List of gene fusions: junction location, gene fusion method used to identify the event including: STARfusion, fusion-CATCHER and DeFUSE (DCC). See Online Methods.

**Supplementary Table 10: Functional enrichment analysis of recurrently altered genes (xls file)**
Enrichment analysis based for lists of genes harboring recurrent SVs. Analysis derived from ToppGene (Online Methods).

### II. SUPPLEMENTARY FIGURES

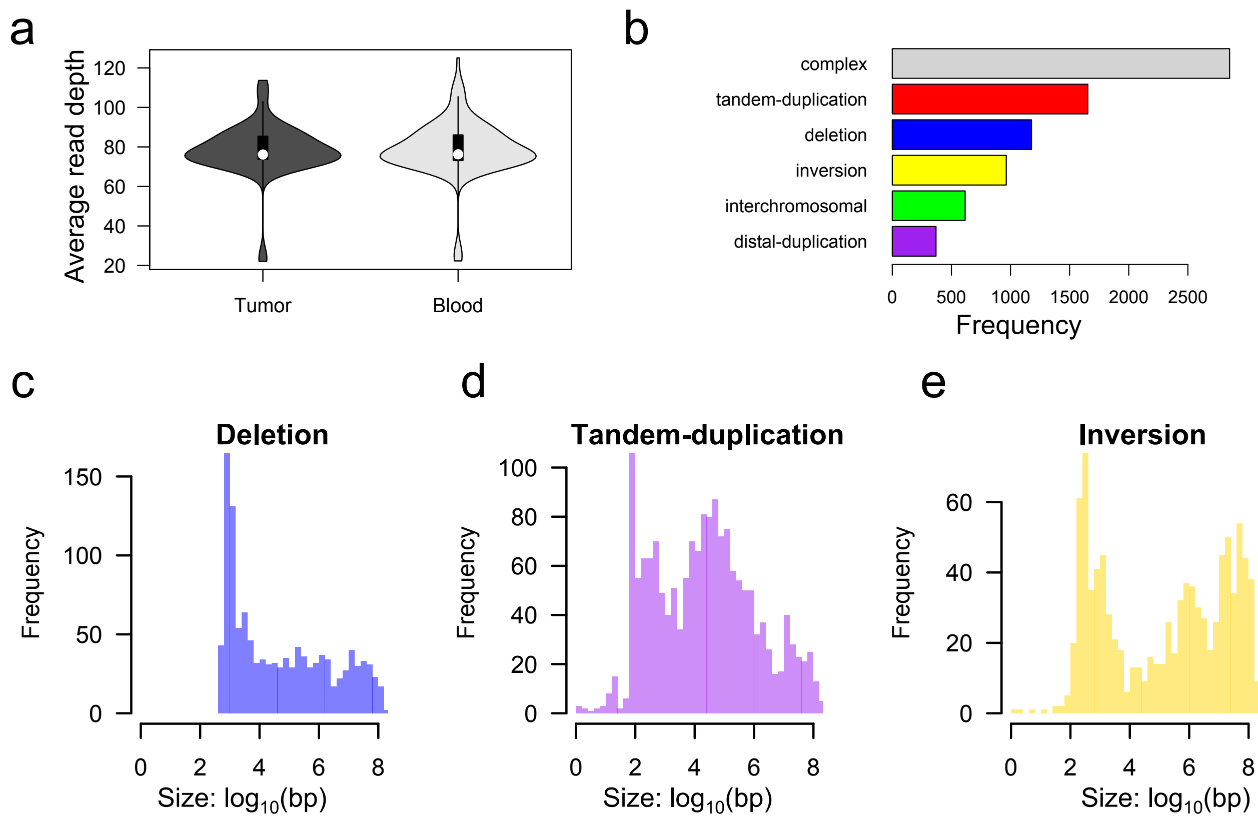

**Supplementary Figure 1: Identification of alignment based structural variants**
**(a)** Average read-depth of primary tumor and matched blood samples. **(b)** Accumulated number of SVs across 135 WGS samples by variant type; inversion bar represents both inversions and probable-inversions **(c-e)** Variant genomic size for **(c)** deletions, **(d)** tandem-duplications and **(e)** inversions + probable-inversion.

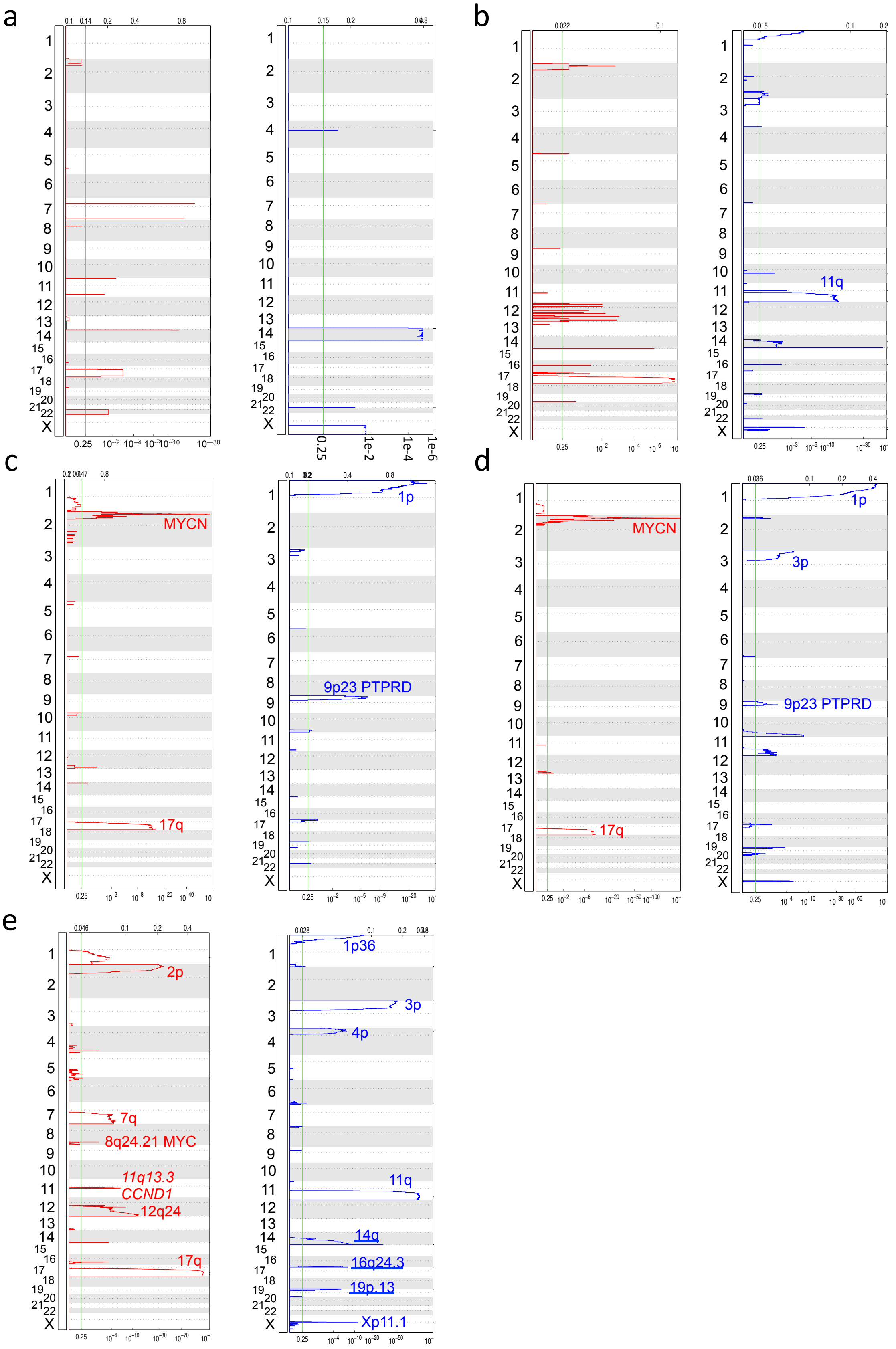

**Supplementary Figure 2: GISTIC analyses for neuroblastomas tumors subtypes**

GISTIC q-plot results for neuroblastomas tumors indicating frequent deletions (blue) and gains (red) obtained from combined LOWR+INTR tumors profiled with **(a)** WGS and **(b)** SNP array. MNA group profiled with **(c)** WGS and **(d)** SNP arrays and **(e)** HR-NA group profiled with SNP arrays.

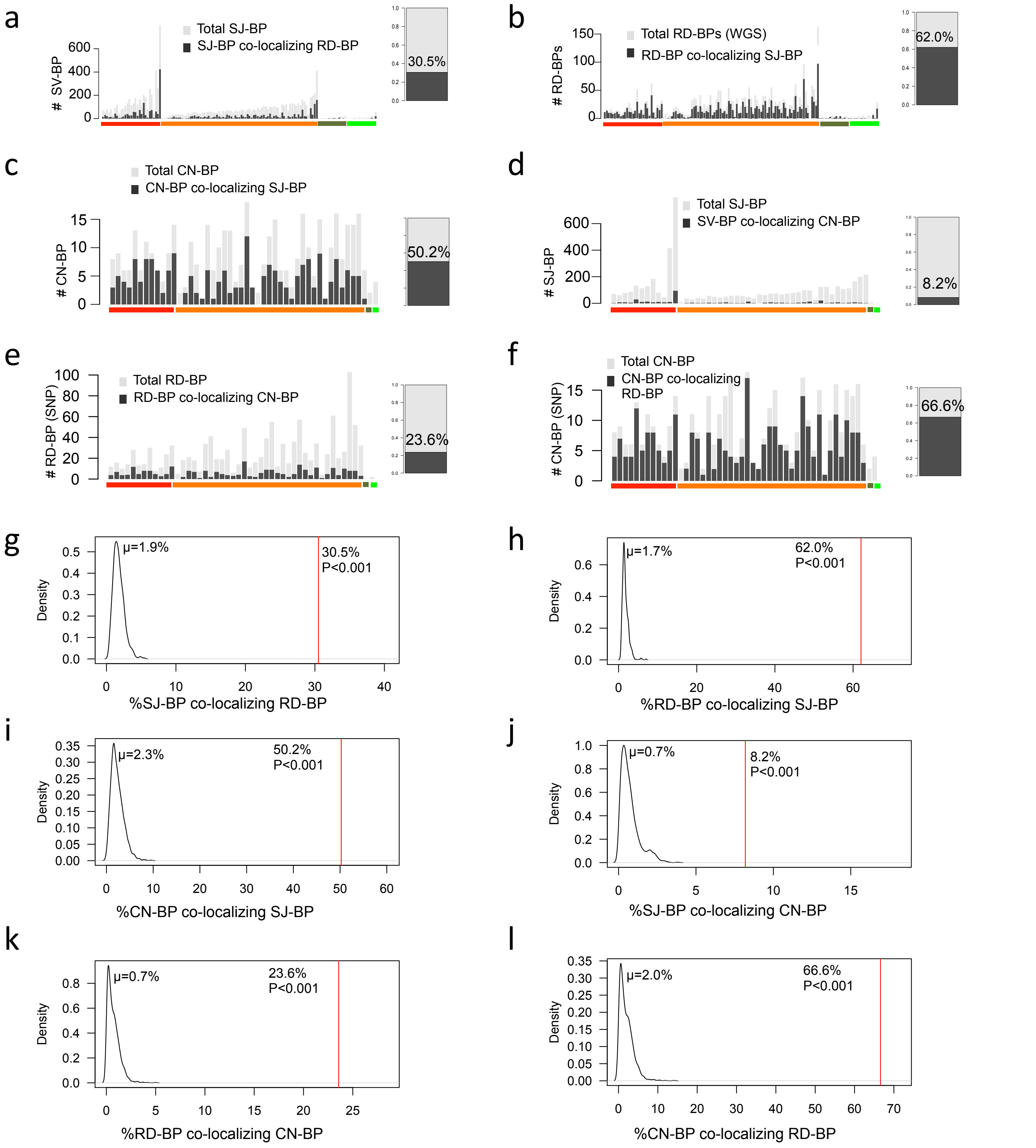

**Supplementary Figure 3: Orthogonal validation of breakpoint detection methods**
**(a-f)** Co-localization of SV breakpoints between alternative SV analyses and platforms. Overall co-localization percentages are located at the right side of each bar plot. **(g-l)** Permutation based randomization of co-localization percentages across breakpoint type comparisons are used to create null distributions and calculate empirical p-values for the observed overall co-localization percentages from (a-f).

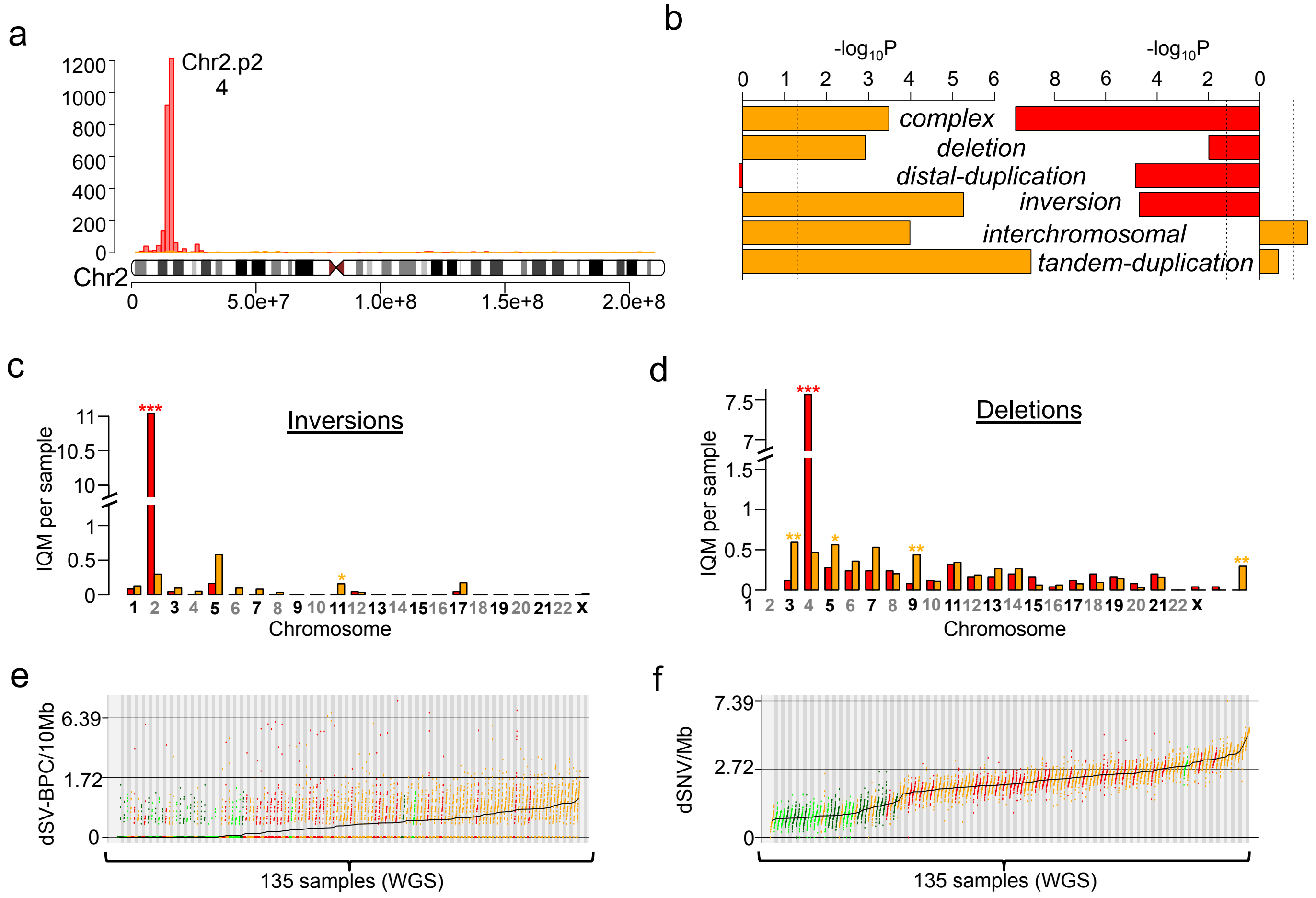

**Supplementary Figure 4: Variability of SV burden across subtypes**

**(a)** Frequency and localization of structural variants in chromosome two across MNA (red) and HR-NA (orange) tumors. **(b)** A comparison of the differential frequencies between MNA and HR-NA of structural variants by type when chromosome 2 is ignored (left) versus included (right); orange bars indicate higher frequency in HR-NA and red bars indicate higher frequencies in MNA; p-values calculated by Wilcoxon rank-sum test. (c-d) By chromosome comparison between MNA and HR-NA of the inter-quantile average number of SVs including **(c)** inversions and **(d)** deletions. **(e-h)** Mutational burden analysis for **(e)** SV and **(f)** SNVs using the IQM burden across 41 chromosome arms.

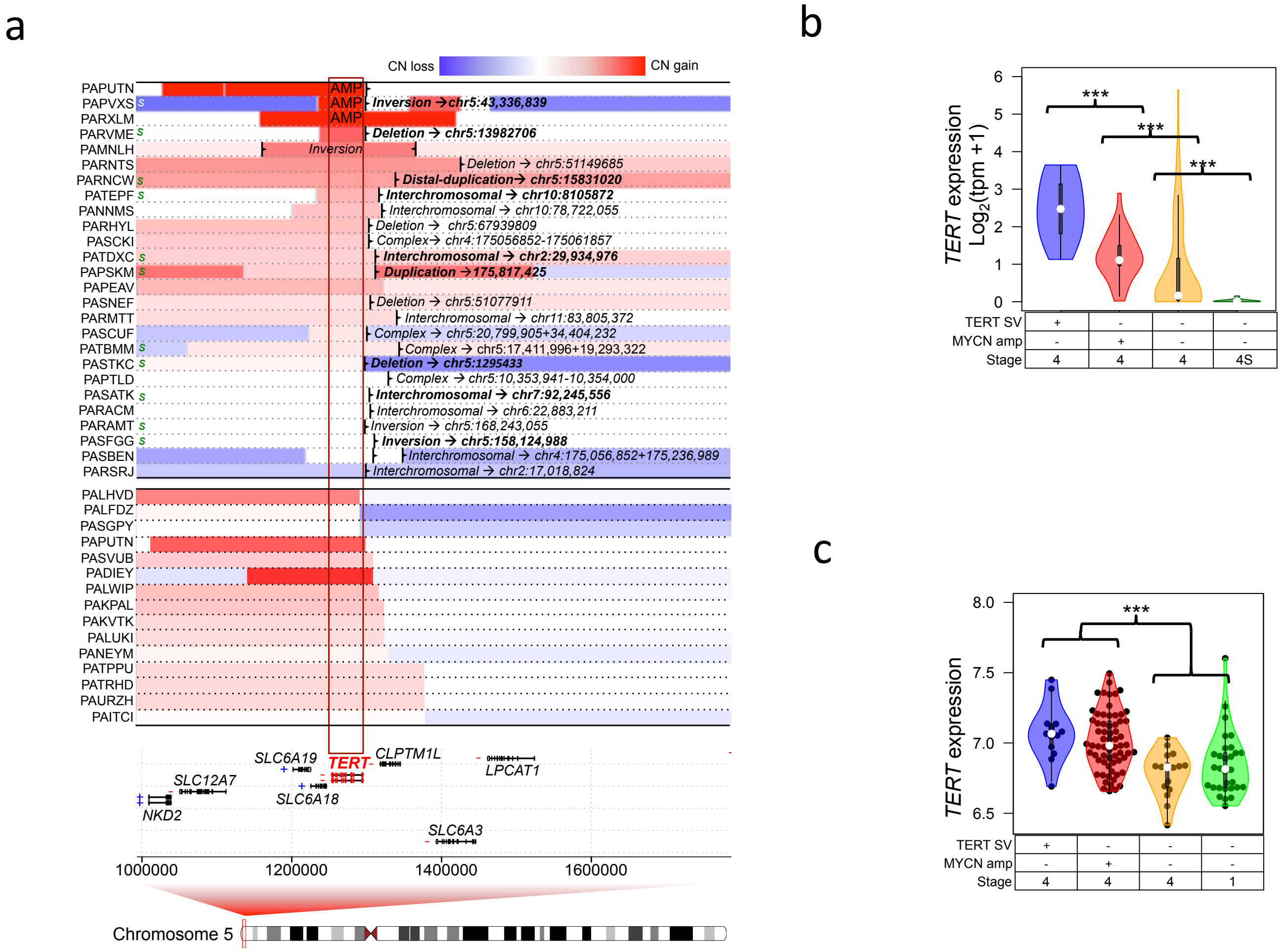

**Supplementary Figure 5: *TERT* rearrangements and expression across neuroblastomas subtypes**
**(a)** SVs at the *TERT* locus in 25 WGS samples based on RD-BPs combined with SJ-BPs and 15 tumors profiled by SNP array using CN-BPs. Nearest upstream *TERT* junctions derived by sequencing are highlighted in text; 'S' at the left of the panel indicates positive validation by Sanger sequencing. **(b)** Expression of *TERT* in primary tumors with different genetic backgrounds obtained from RNA-seq TARGET cohort. **(c)** Expression of *TERT* in primary tumors with different genetic backgrounds obtained from HumanExon arrays TARGET cohort.

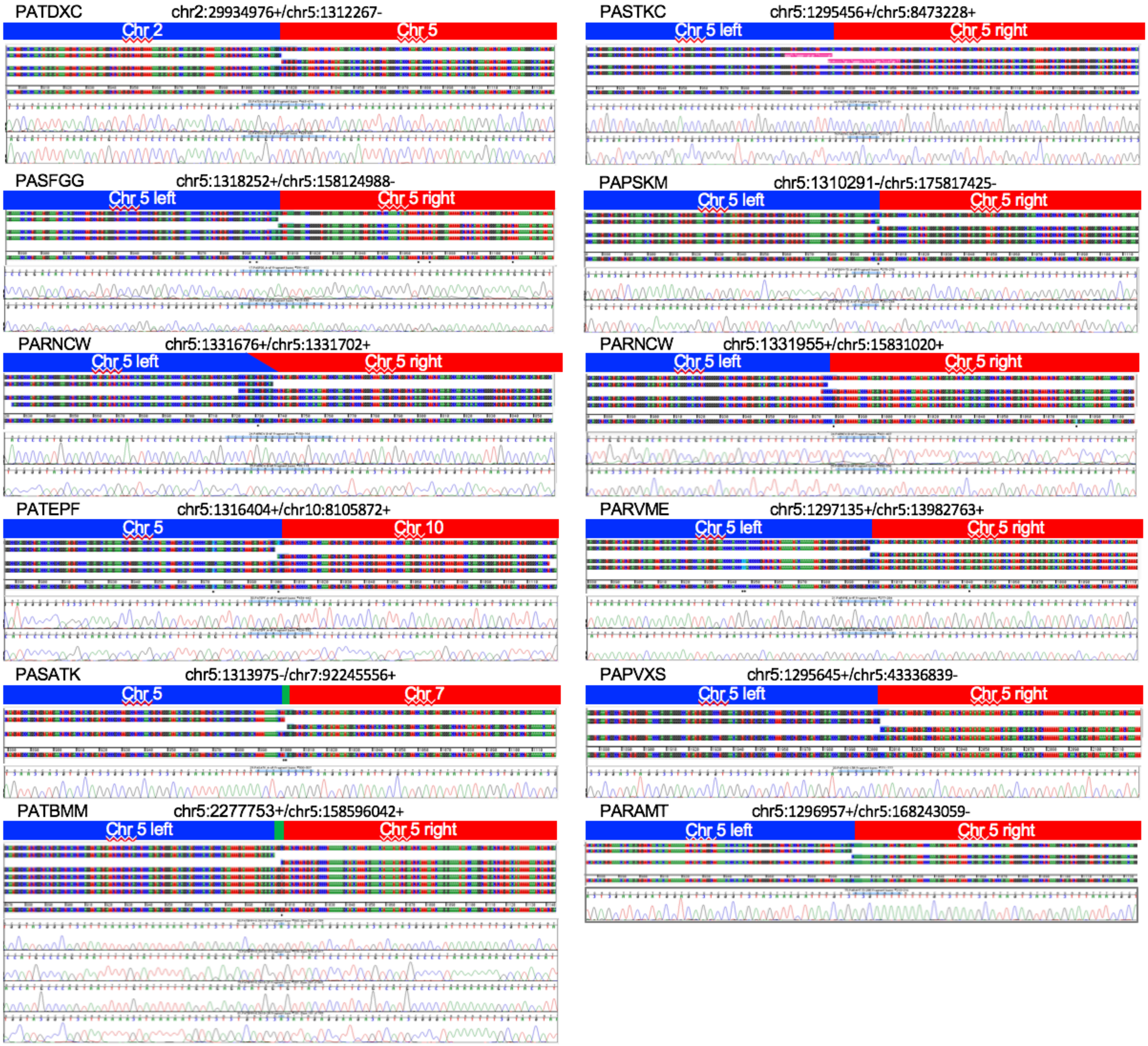

**Supplementary Figure 6: Sanger sequencing validation of rearrangements near *TERT* gene**
Primers and additional details described in Supplementary Table 8.

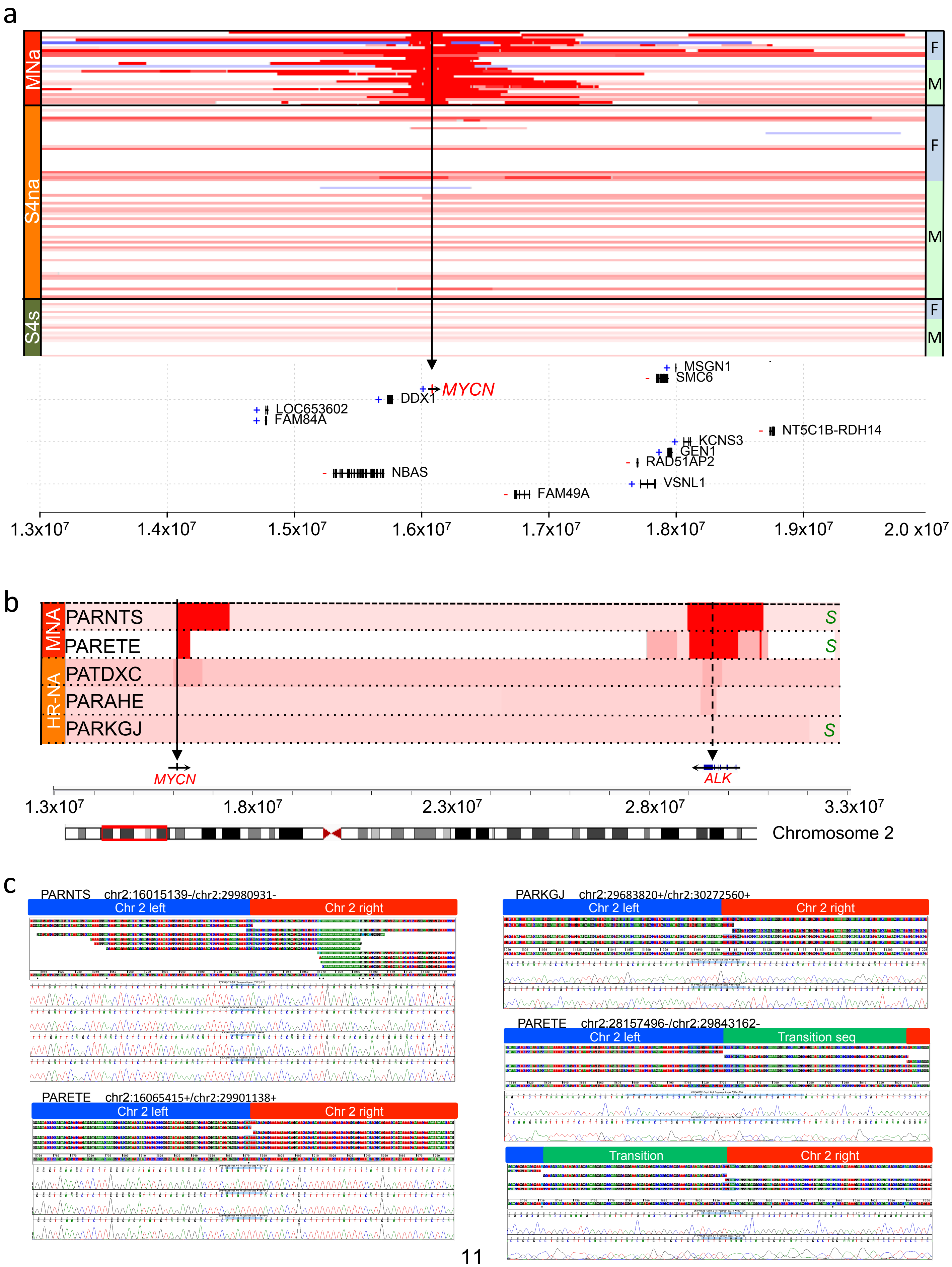
a
MNa
S4na
S4s
F
M
MYCN
DDX1
LOC653602
FAM84A
NBAS
MSGN1
SMC6
KCNS3
GEN1
RAD51AP2
VSNL1
FAM49A
NT5C1B-RDH14
1.3x10^7
1.4x10^7
1.5x10^7
1.6x10^7
1.7x10^7
1.8x10^7
1.9x10^7
2.0 x10^7
b
MNA
HR-NA
PARNTS
PARETE
PATDXC
PARAHE
PARKGJ
S
MYCN
ALK
1.3x10^7
1.8x10^7
2.3x10^7
2.8x10^7
3.3x10^7
Chromosome 2
c
PARNTS chr2:16015139-/chr2:29980931-
Chr 2 left
Chr 2 right
PARETE chr2:16065415+/chr2:29901138+
Chr 2 left
Chr 2 right
PARKGJ chr2:29683820+/chr2:30272560+
Chr 2 left
Chr 2 right
PARETE chr2:28157496-/chr2:29843162-
Chr 2 left
Transition seq
Transition
Chr 2 right

**Supplementary Figure 7: *MYCN* and *ALK* structural variants and Sanger validation**

**(a)** IGV track visualization plot shows copy number segmentation data around the MYCN amplicon (chr2p.24) region. Three groups separated at the left margin: MNA (red), HR-NA (orange) and S4s (dark green). Left margin separates samples by gender within each tumor group: female (light blue) and male (light green). **(b)** IGV track visualization plot shows copy number segmentation data comprising MYCN and ALK loci in cases with ALK associated SVs; variant position and type highlighted for ALK; “S” at the left of the panel indicates that Sanger sequencing validation is availabe. **(c)** Sanger sequencing validation of rearrangements near *ALK* gene in 3 cases with available DNA (PARETE harbored multiple SVs). Primers and additional details described in Supplementary Table 8.

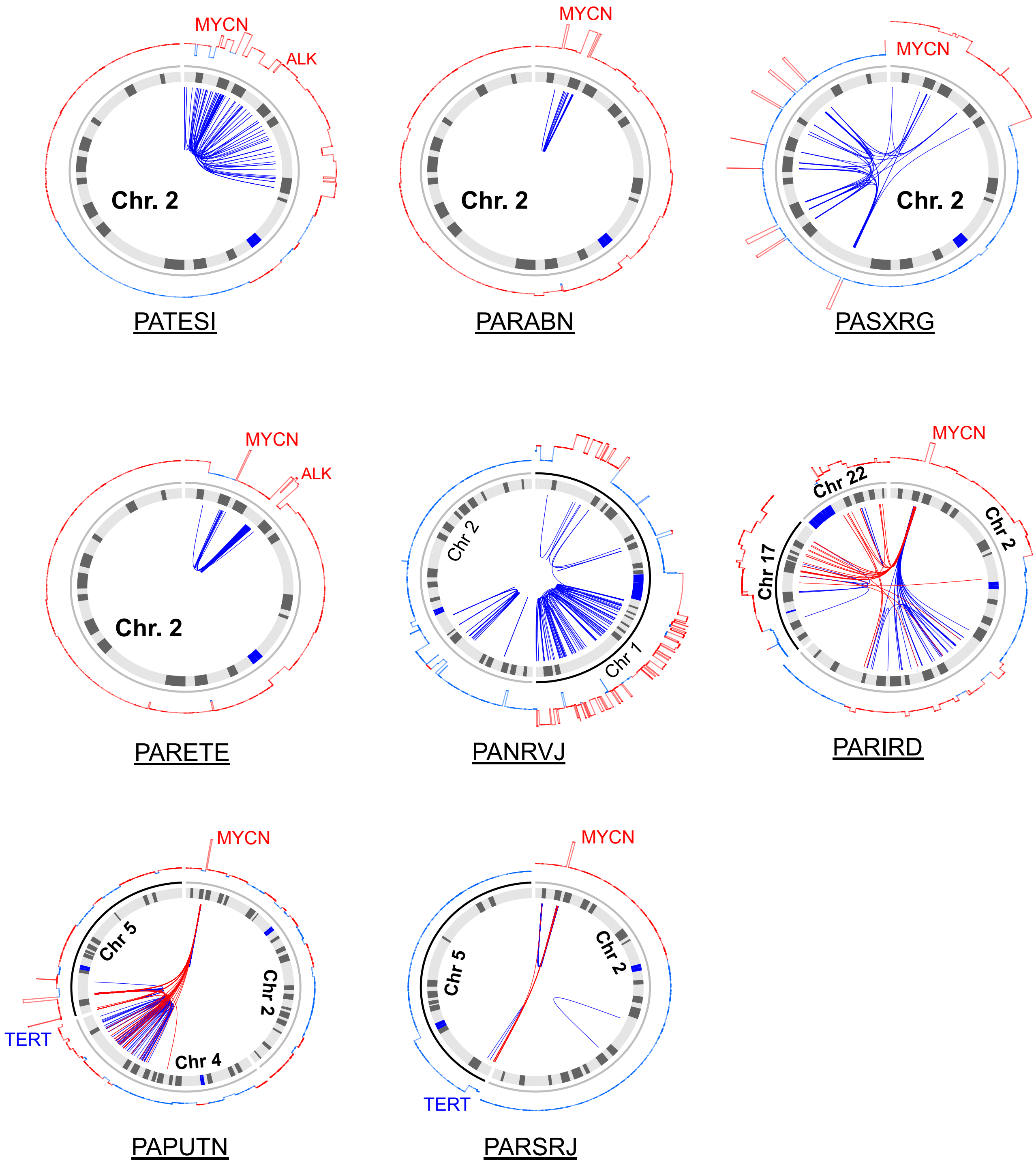

**Supplementary Figure 8: Chromothripsis in chromosome 2 associates with *MYCN* amplification**
Circos plots representing copy number and large structural variants (<100Kb) in samples with high BP and SV density indicative of chromothripsis. Samples with MYCN amplification are highlighted in red text.

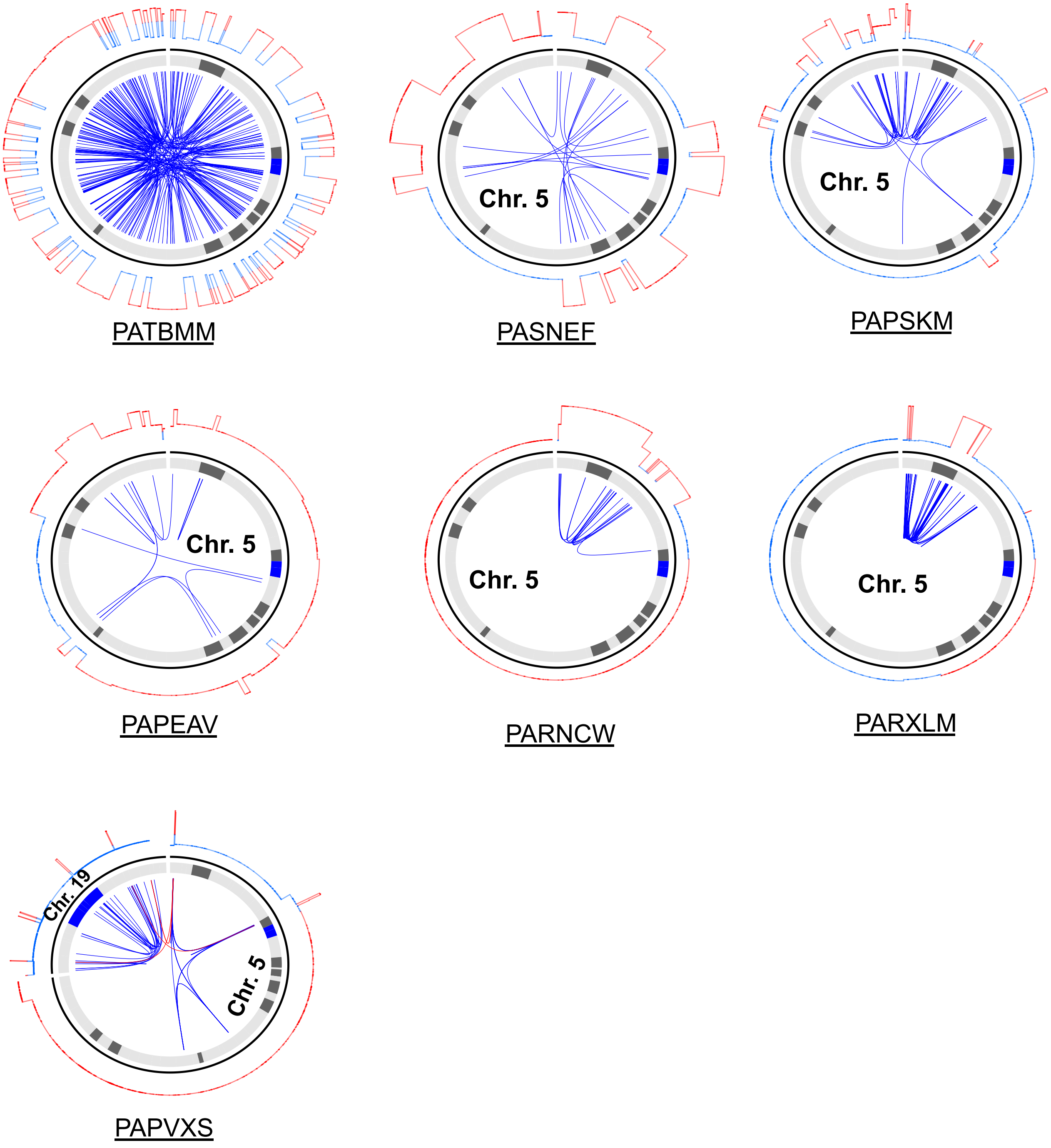

**Supplementary Figure 9: Chromothripsis on chromosome 5 associates with *TERT* rearrangements**
Circos plots representing copy number and large structural variants (<100Kb) in samples with high BP and SV density indicative of chromothripsis as derived from Supplementary Table 3.

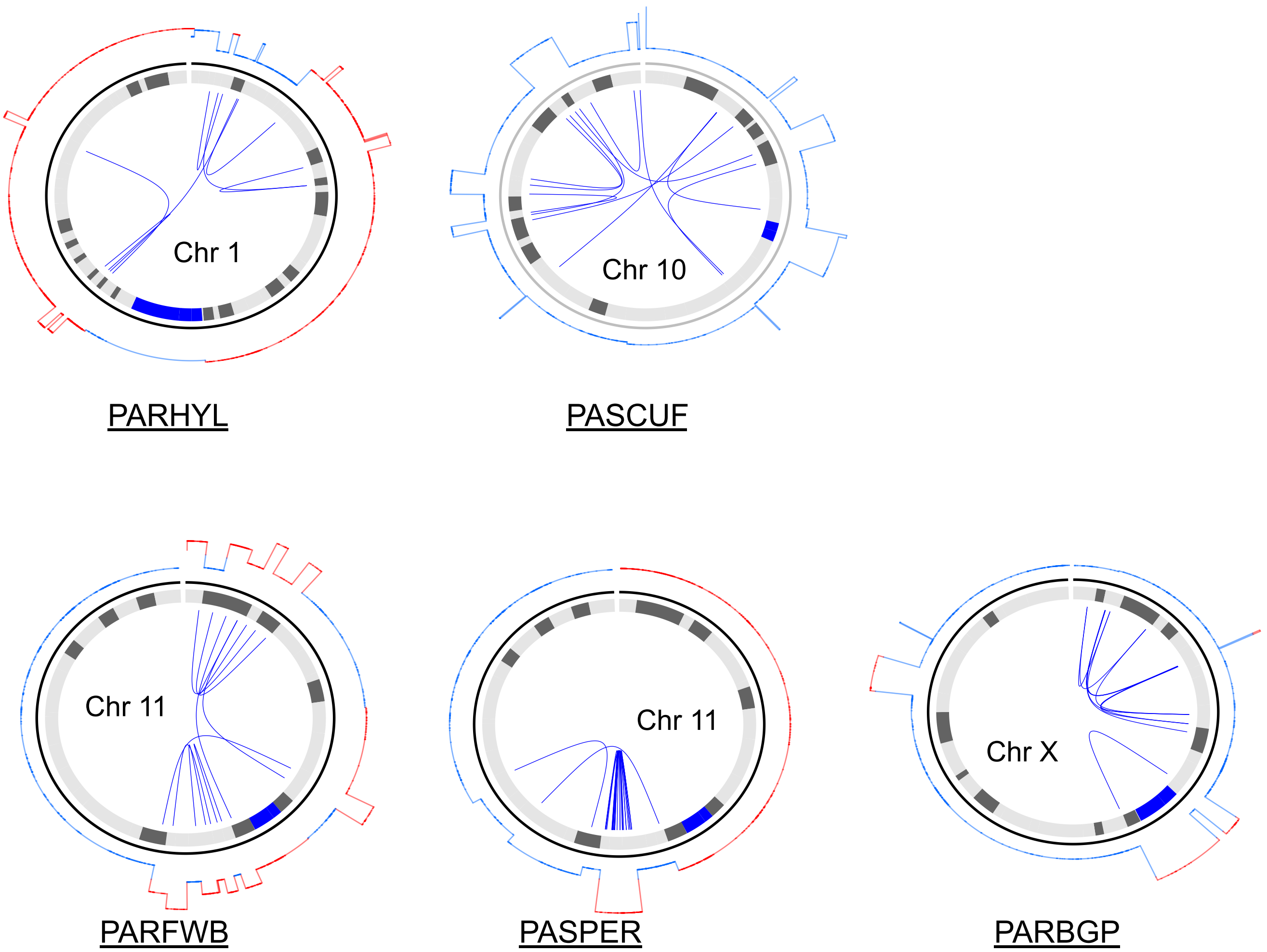

**Supplementary Figure 10: Chromothripsis in chromosomes 1, 10, 11 and X**
Circos plots representing copy number and large structural variants (<100Kb) in samples with high BP and SV density indicative of chromothripsis as derived from Supplementary Table 3.

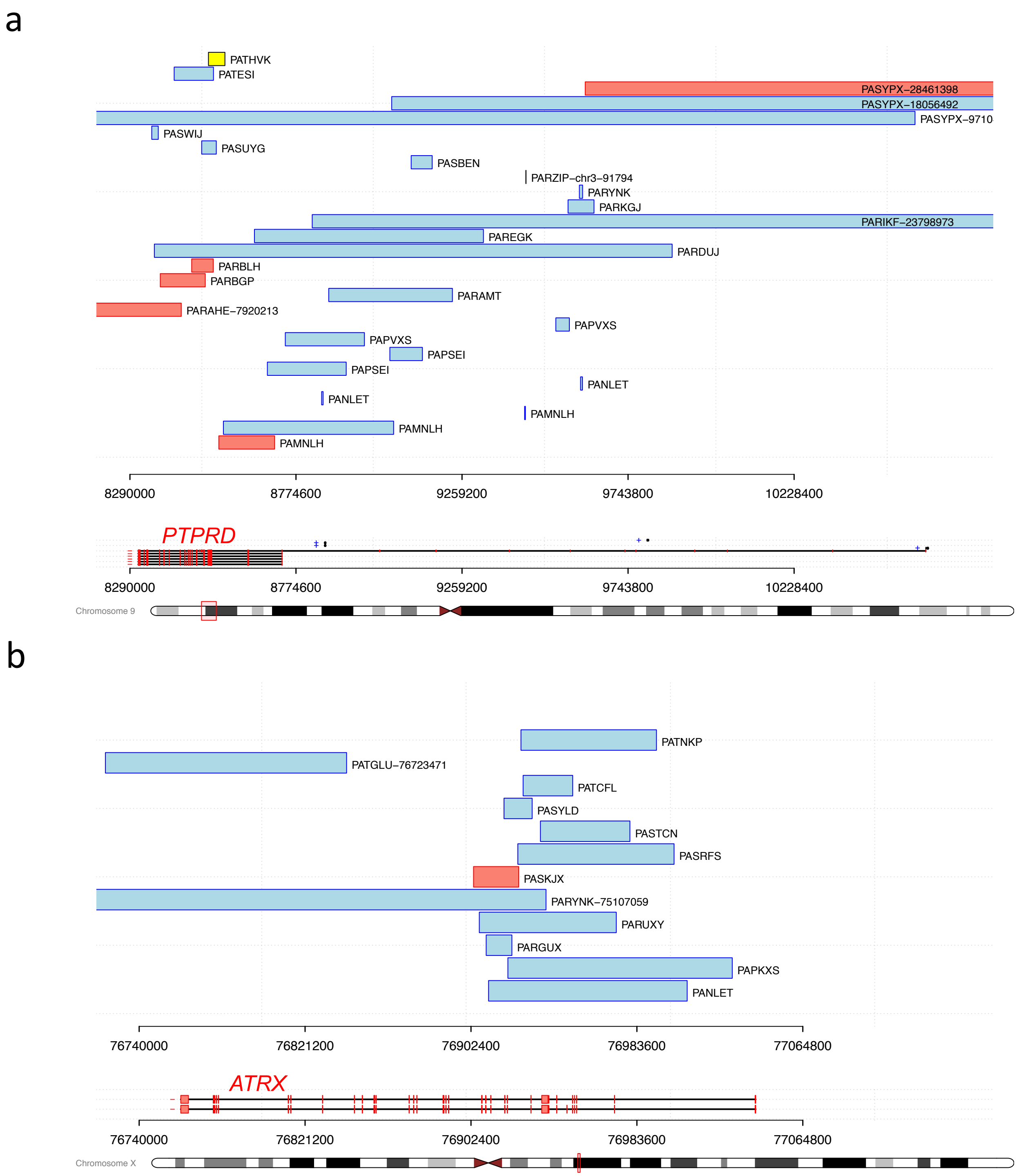

**Supplementary Figure 11: Recurrently altered genes: *PTPRD* and *ATRX***
Genomic visualization of sequence junction based structural variants in **(a)** *PTPRD* and **(b)** *ATRX* loci. Boxes represent the spanned region of deletions (blue), duplications (red) and inversions (yellow). Translocations are represented as vertical line followed by the sample names and genomic coordinate of the destination region. Coordinates of structural variants outside the plot genomic span also included in the labels.

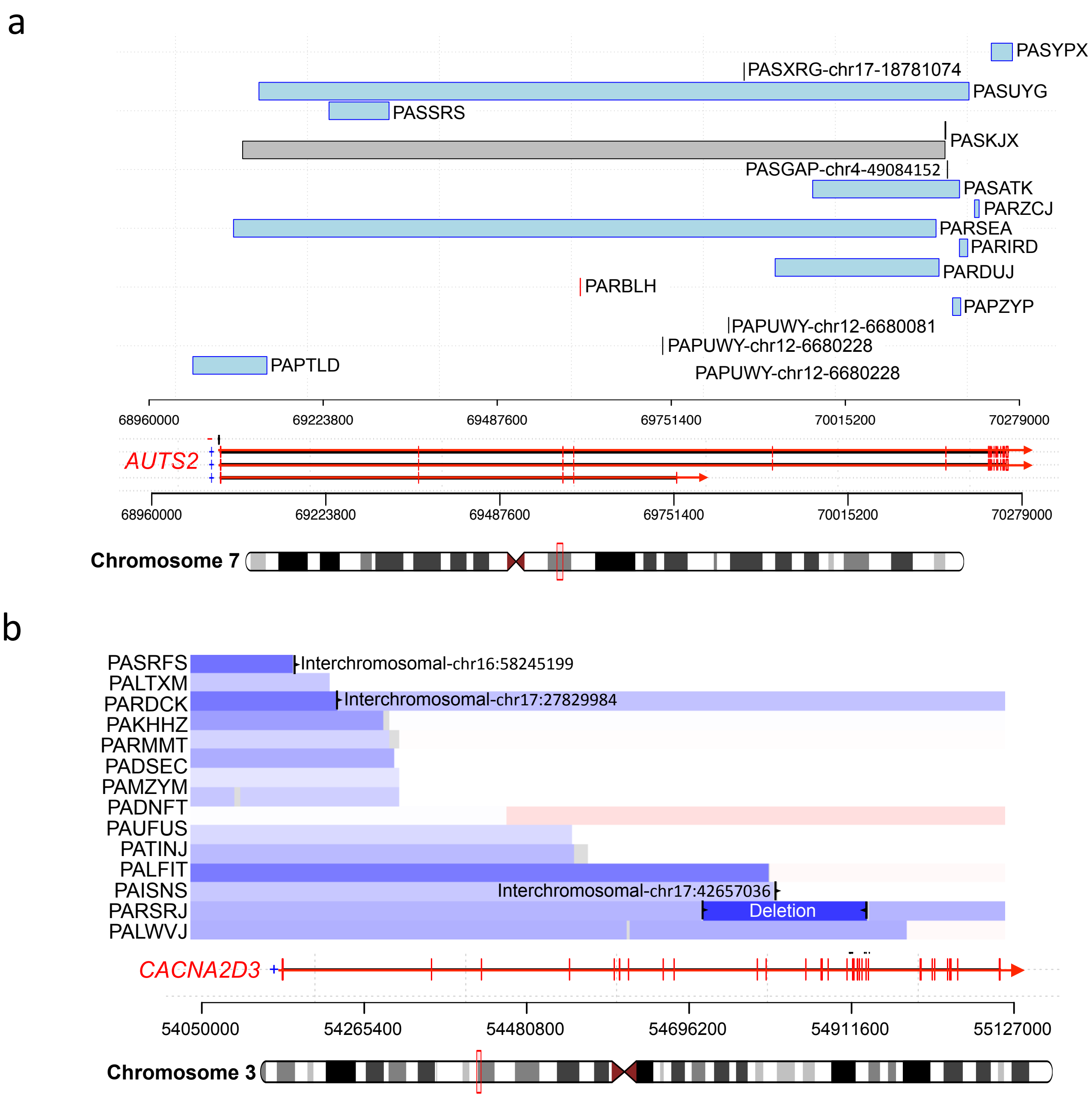

**Supplementary Figure 12: Recurrently altered genes: *AUTS2* and *CACNA2D3***

**(a)** Sequence junction based structural variants in *AUTS2 locus.* Boxes represent the spanned region of deletions (blue) and duplications (red) and inversions (yellow). Translocations are represented as vertical line followed by the sample names and genomic coordinate of the destination region. Coordinates of structural variants outside the plot genomic span also included in the labels. **(b)** IGV copy number visualization of *CACNA2D3* locus with sequence junctions overlay.

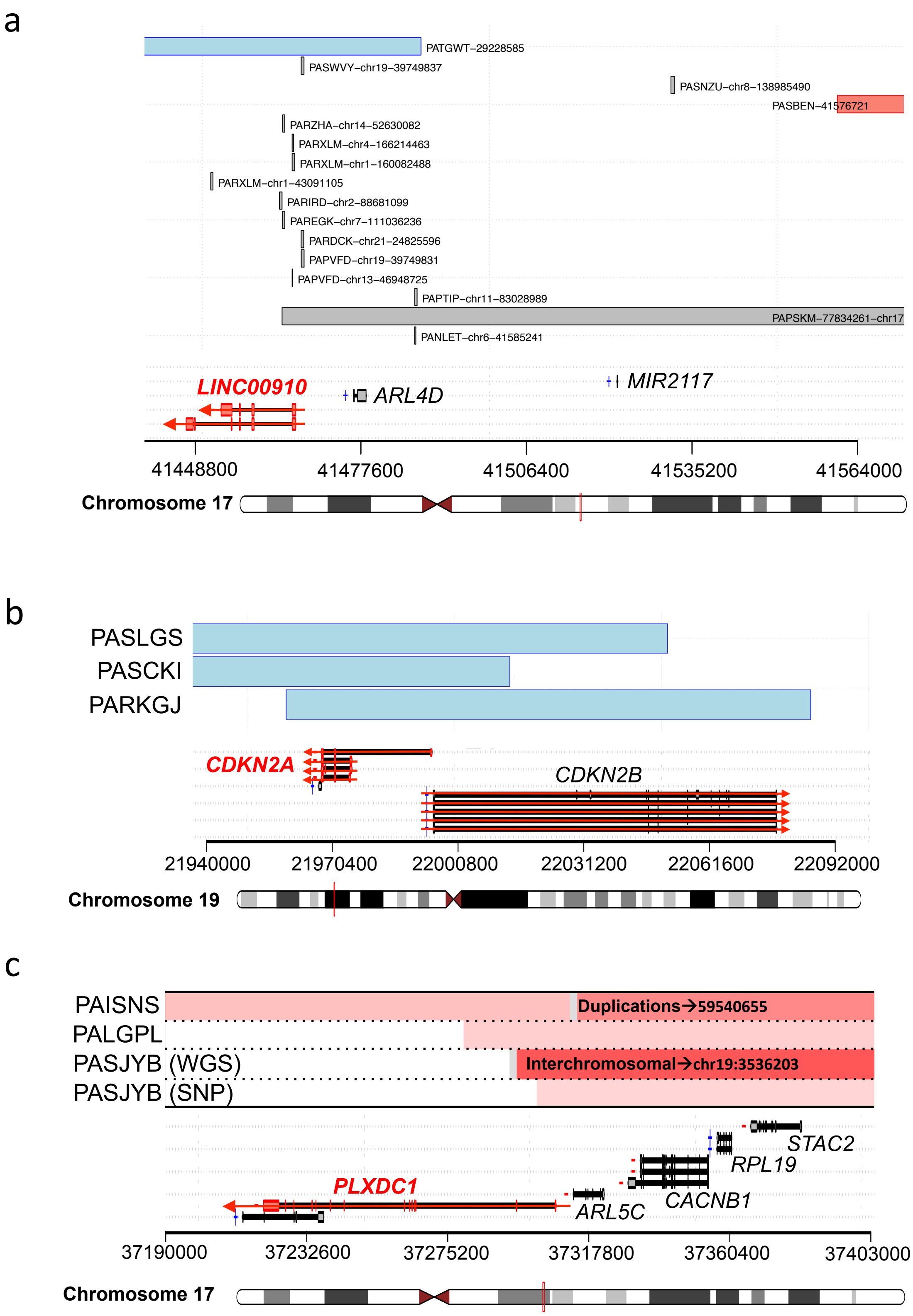

**Supplementary Figure 13: Recurrently altered genes: *LINC00910*, *CDKN2A* and *PLXDC***
Genomic visualization of breakpoints implicated in eQTL associations of **(a)** *LINC00910,* **(b)** *CDKN2A* and **(c)** *PLXDC1 loci*. Information was derived from sequence junctions in **(a)** and read-depth copy number for **(b)** and **(c).** Regions in blue indicate deletions or copy number loss, regions in red indicate duplications and gains. Translocations are represented as vertical line followed by the sample names and genomic coordinate of the destination region.

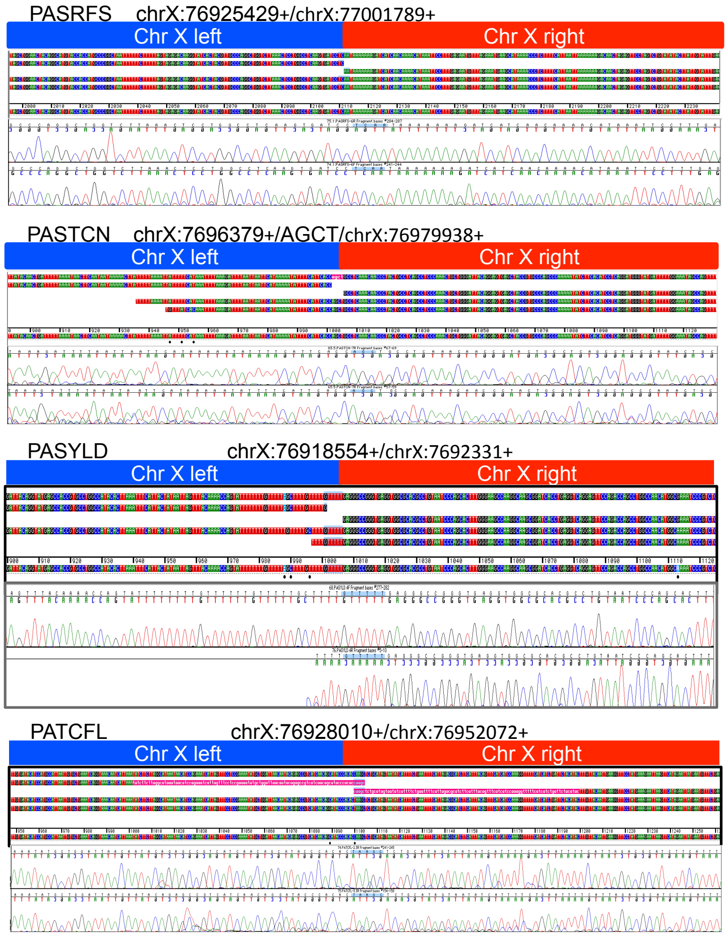

**Supplementary Figure 14: Sanger sequencing validation of *ATRX* deletions**
Primers and additional details described in Supplementary Table 8.

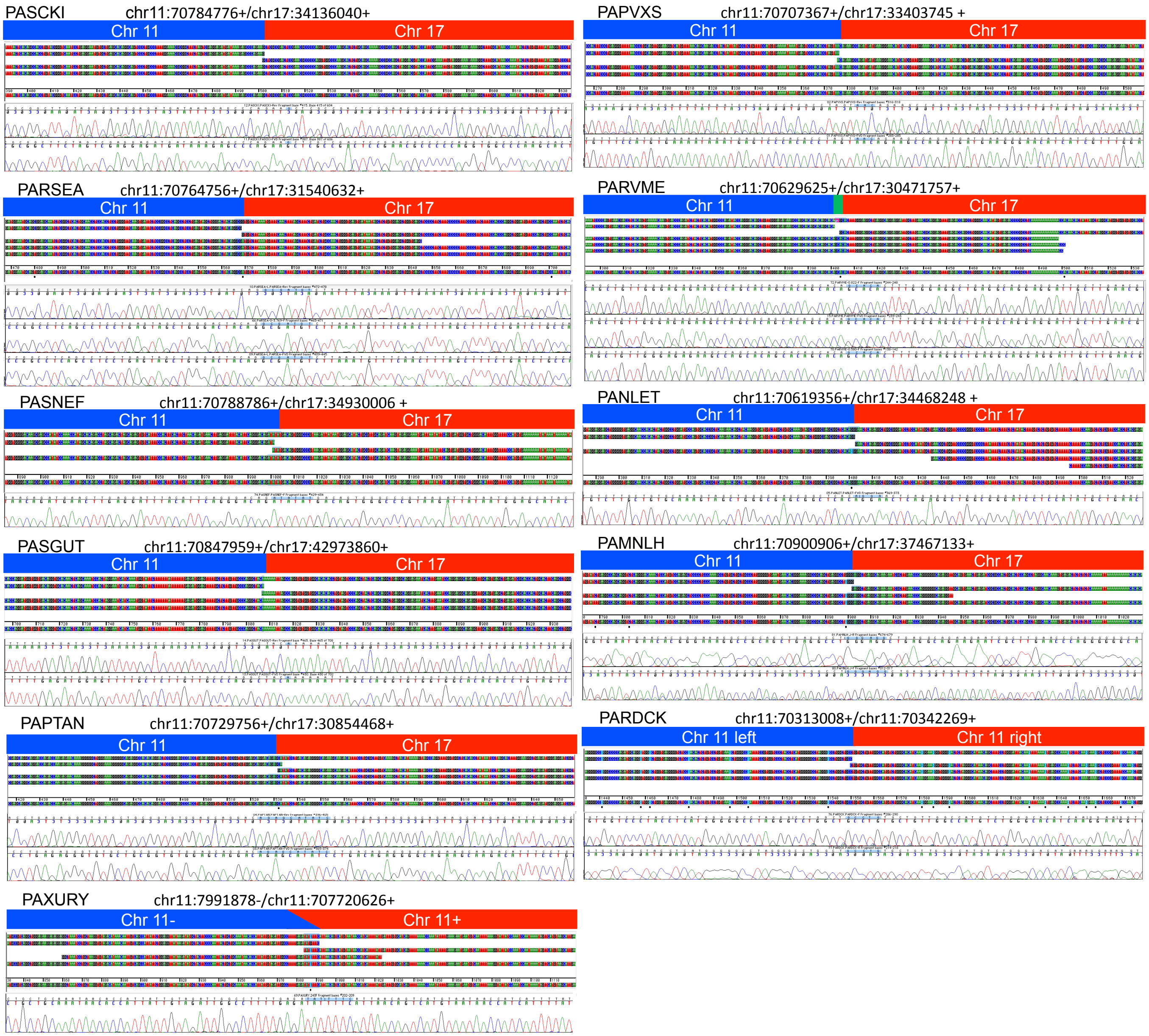

**Supplementary Figure 15: Sanger sequencing validation of *SHANK2* structural variants**
Primers and additional details described in Supplementary Table 8.

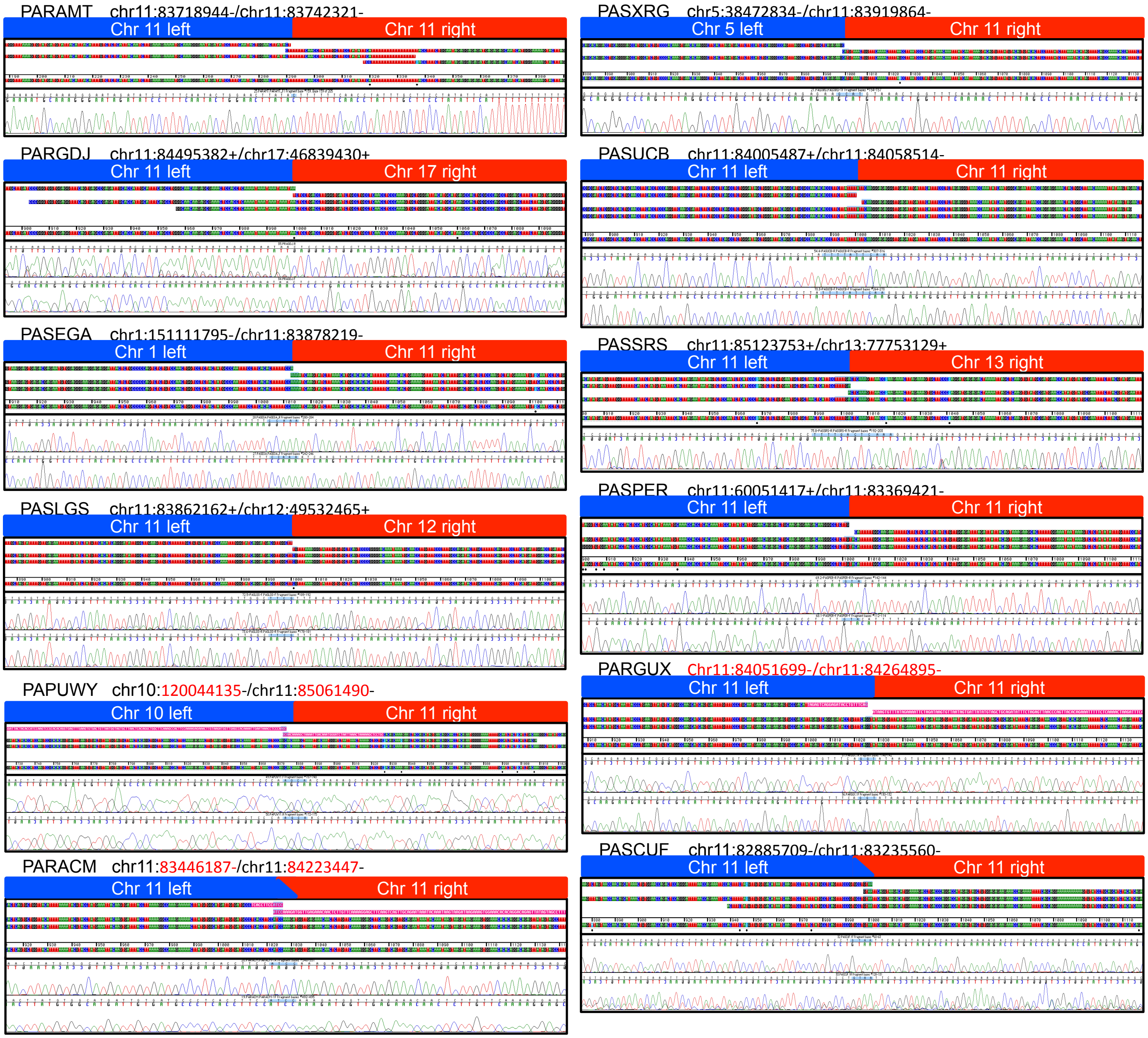

**Supplementary Figure 16: Sanger sequencing validation of DLG2 rearrangements**
Primers and additional details described in Supplementary Table 8.

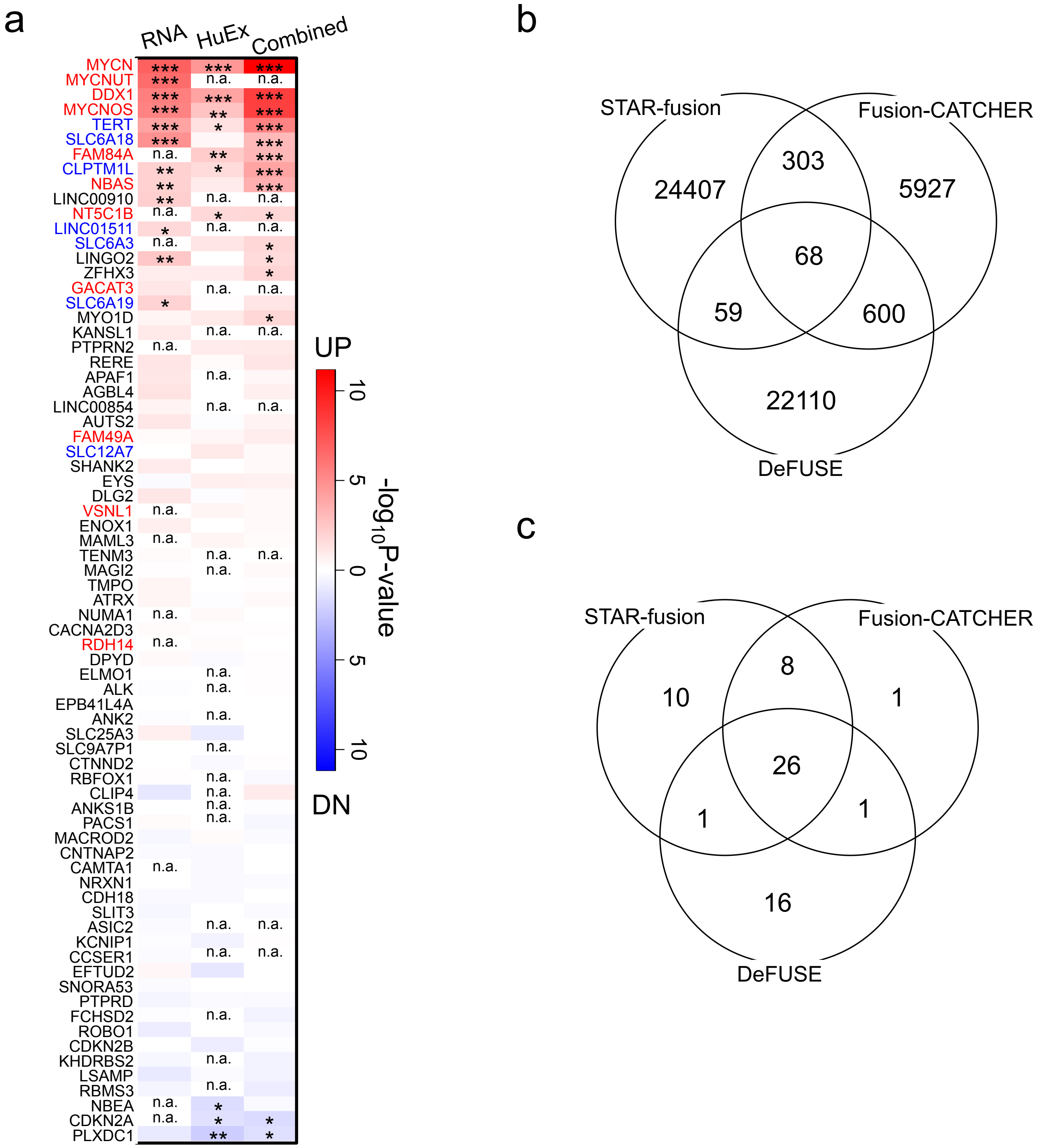

**Supplementary Figure 17: Transcriptional effect of structural variants: eQTL and gene fusions**
**(a)** Differential expression analysis of genes with samples harboring SVs compared to unaltered samples using a Wilcoxon rank sum test. Gene names colored in red map into Chr2.p24 region near *MYCN*; Genes colored in blue map into Chr5.p15 near *TERT*. Asterisk indicates the p-value cut off (*** = P < 0.001; ** = P<0.01; *= P<0.05); n.a. indicates that the test couldn't be done due to lack of data points or because the gene was not included in the HumanExon array probeset design. **(b)** Overlap across three different gene fusion callers from 153 RNA-seq neuroblastoma samples. **(c)** Overlap across three different gene fusion callers from RNA-seq with evidence of translocation events in 89 samples with both RNA-seq and WGS profiles, see also Supplementary Table 9.

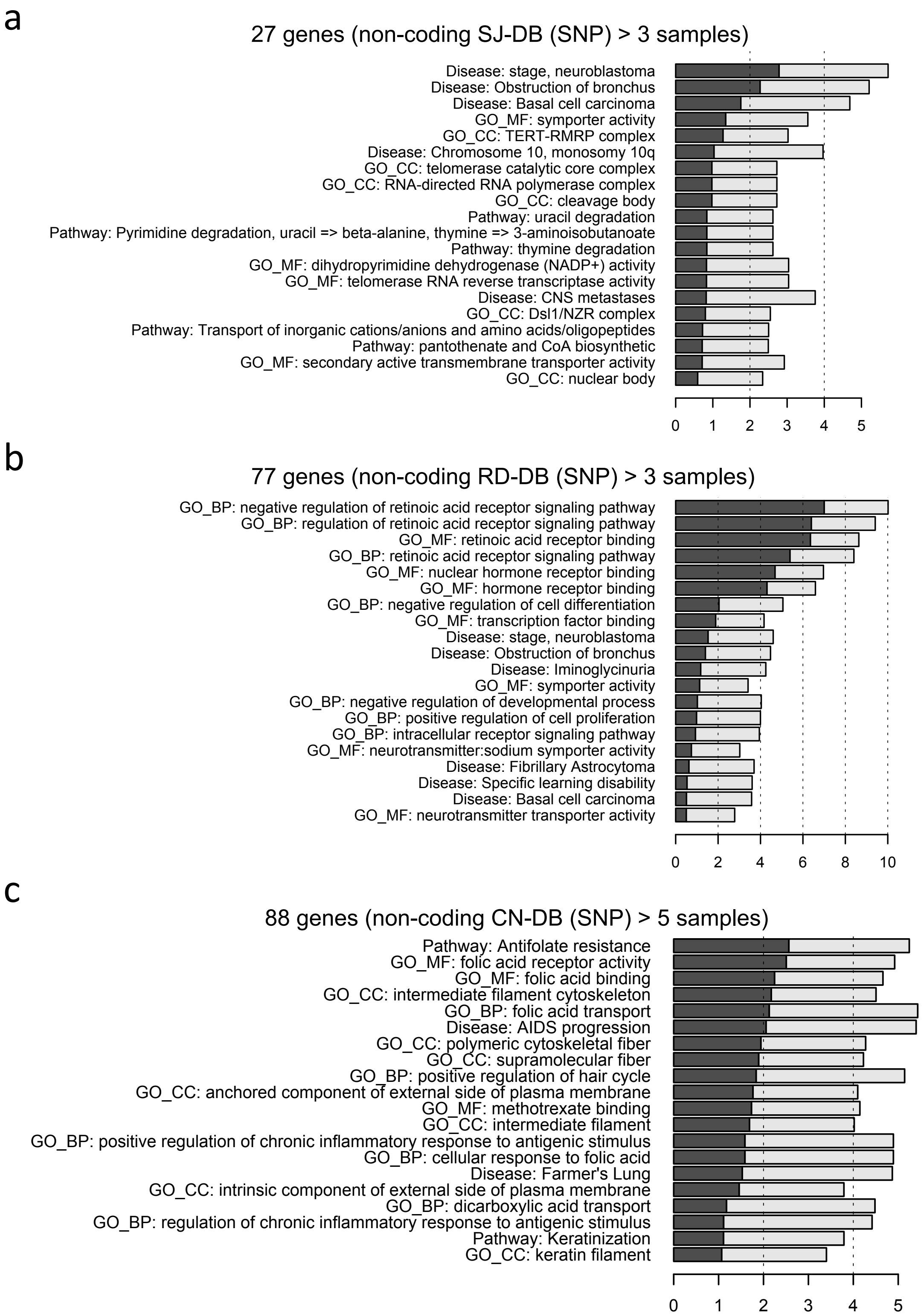

**Supplementary Figure 18: Functional enrichment of genes with proximal and intronic variants**
**(a-c)** Function enrichment analysis bar plots for genes recurrently altered based on proximal and intronic breakpoint analyses of **(a)** SJ-BPs, **(b)** RD-BPs and **(c)** CN-BPs.

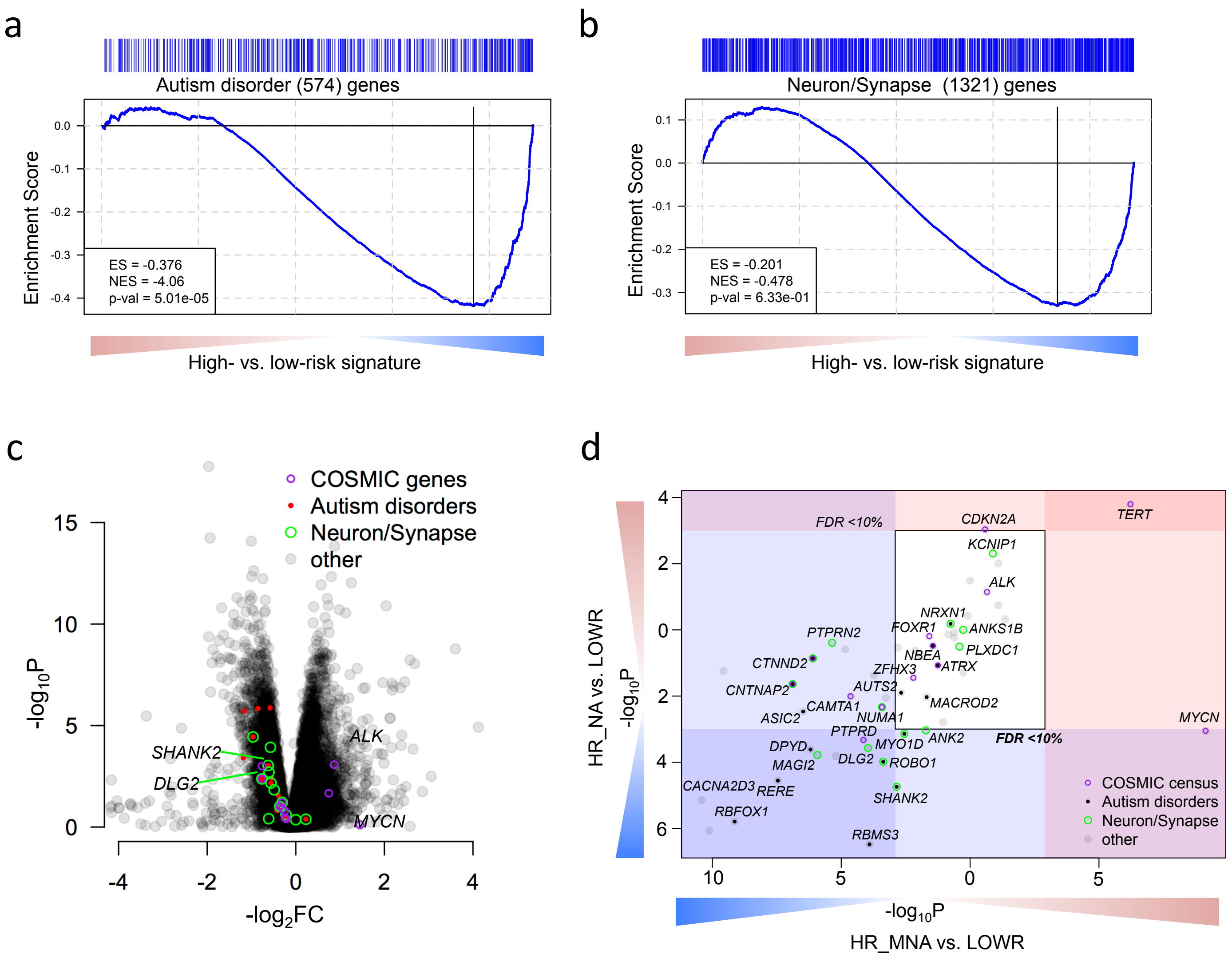

**Supplementary Figure 19: Neurodevelopmental pathways are down-regulated in high-risk neuroblastoma**

**(a-b)** Gene Set Enrichment Analysis across the signature of high- versus low-risk tumors from the HumanExon array show enrichment of **(a)** autism spectrum disorder predisposition genes and **(b)** neuronal and synapse part genes. **(c)** Volcano plot showing differential expression between high- and low-risk highlighting genes with recurrent SVs and their functional classification **(d)** Subtype specific high- versus low-risk differential expression analysis of 77 recurrently altered genes from Fig 4i shown as scatter plot (MNA = x-axis, HR-NA = y axis). (d-g) Analysis replicated in two datasets: RNA-seq (here) and HuEx arrays (Fig. 5 c-f).

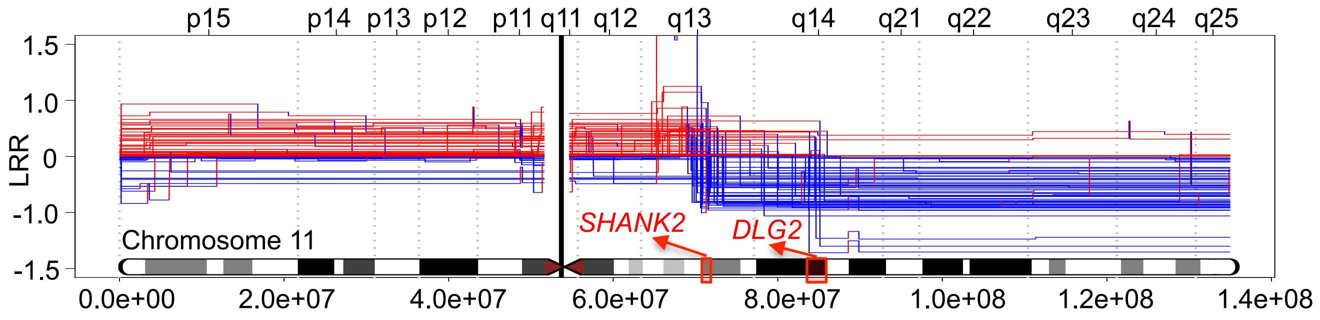

**Supplementary Figure 20: Chromosome 11 breakpoints frequently map into *SHANK2* and *DLG2 loci***
Chromosome 11 segmentation plot for HR-NA tumors obtained from WGS highlights regions of high breakpoint density near *SHANK2* and *DLG2* loci.

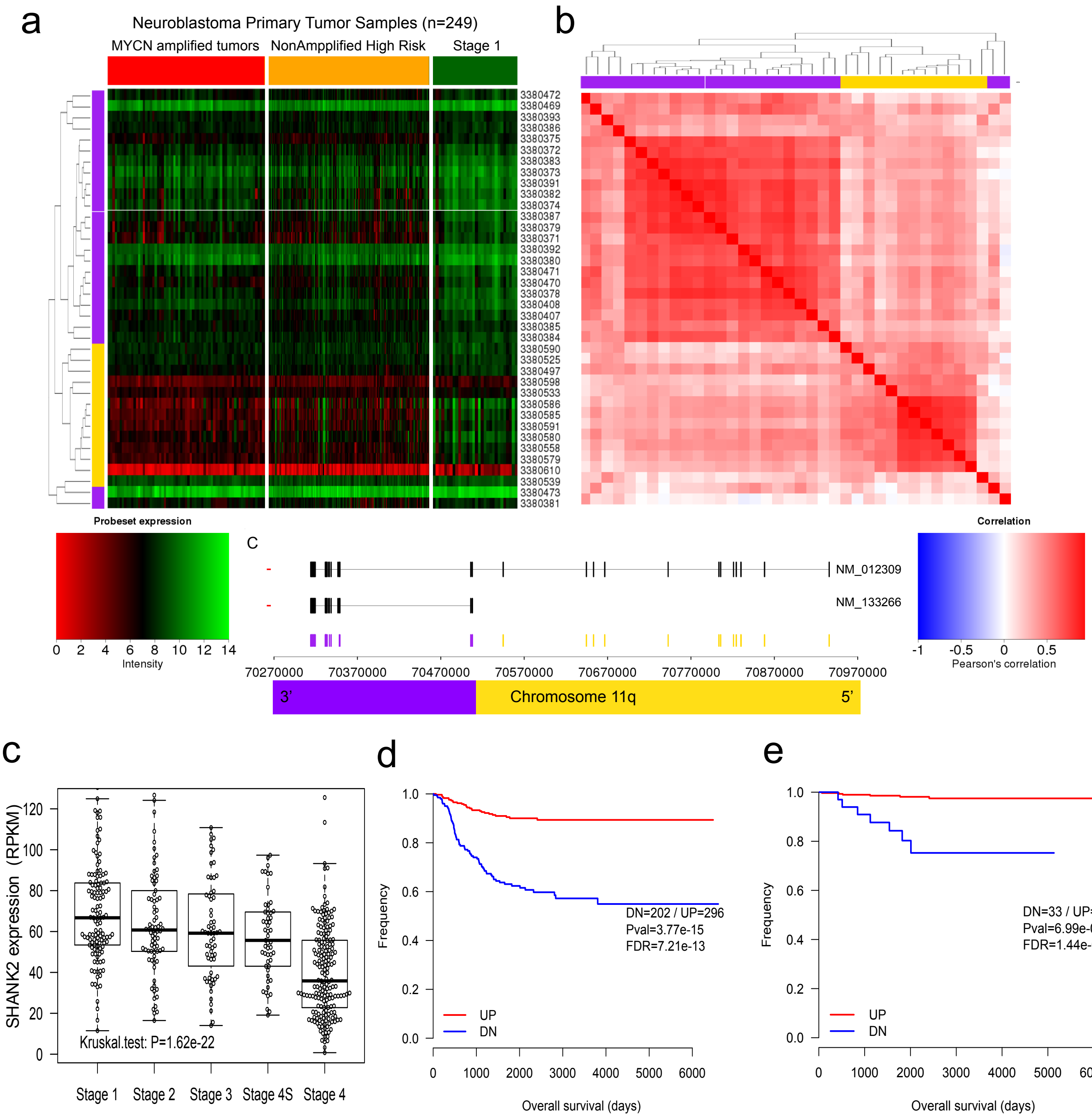

**Supplementary Figure 21: Low *SHANK2* expression in neuroblastoma is associated with poor survival. (a,b)** Clustering analysis of *SHANK2* exon level expression from Affymetrix HumanExon arrays in MNa, S4na and S1 tumors. The heatmap **(a)** shows higher exon expression level in S1 compared to MNa and S4na. The correlation matrix **(b)** shows two well-defined clusters associated with the two known coding isoforms of the gene. **(c)** *SHANK2* long isoform (NM_012309) expression decreases in high INSS stage across 498 primary neuroblastomas (SEQC dataset). **(d)** Kaplan-Meier analysis of *SHANK2* long isoform (NM_012309) expression shows association of low expression with poor outcome (SEQC dataset, all samples). **(e)** decreased *SHANK2* expression (NM_012309) is associated with reduced overall survival within the low- and intermediate-risk subsets (excluding high-risk).

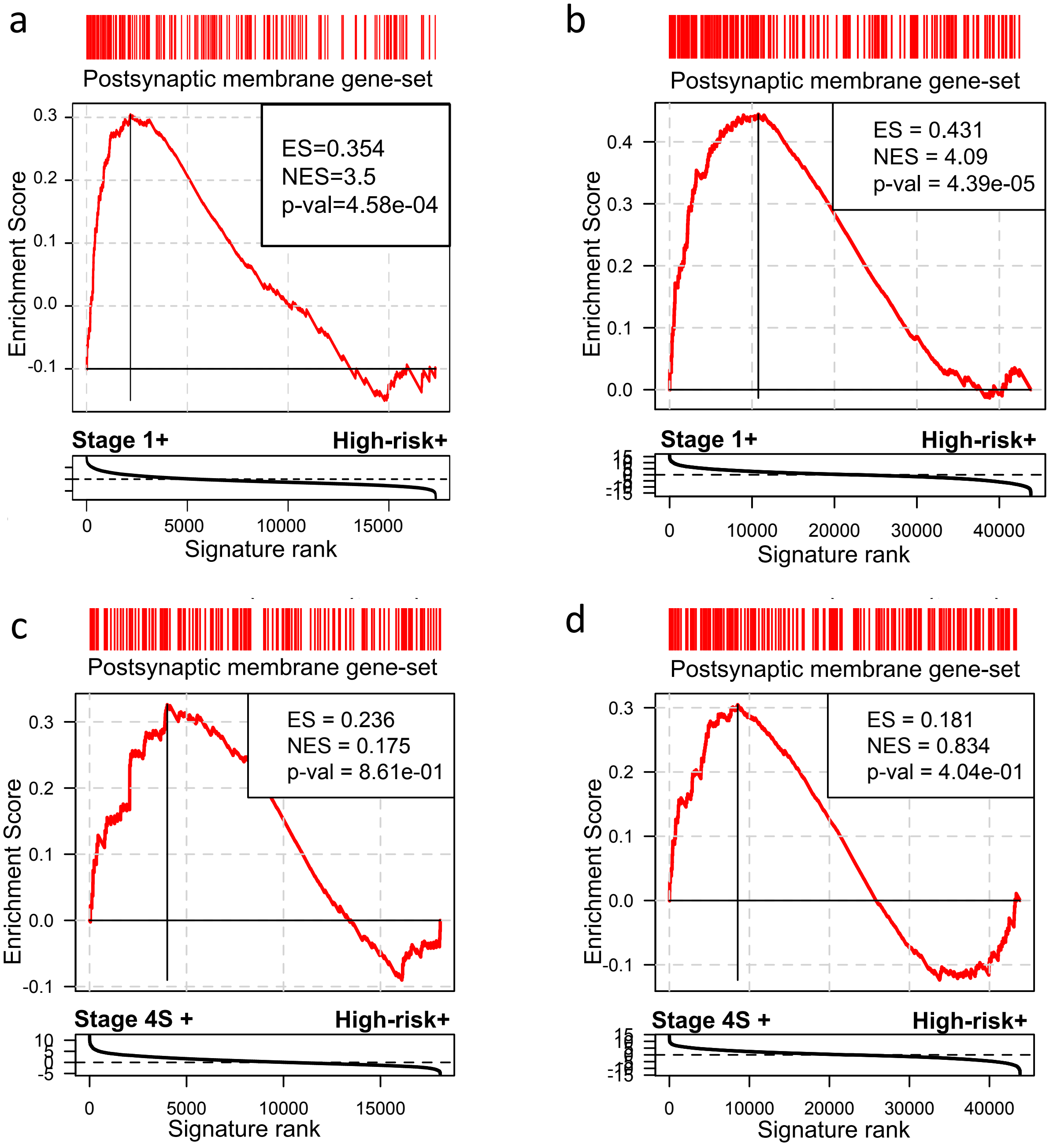

**Supplementary Figure 22: Genes of the postsynaptic density are down-regulated in high-risk neuroblastoma**

"Postsynaptic membrane" genes were obtained from MsigDB Gene Ontology Cellular Component gene set library and subjected to GSEA analysis of differential gene expression (DGE) signatures. DGE were obtained by comparing high-risk tumor's expression against stage 1 and 4S in different datasets: **(a)** TARGET Affymetrix Human Exon Stage 1 vs. High-Risk, **(b)** SEQC 498 RNA-seq Stage 1 vs. High-risk, **(c)** TARGET RNA-seq Stage 4S vs. High-risk and **(d)** SEQC 498 RNA-seq Stage 4S vs. High-risk.

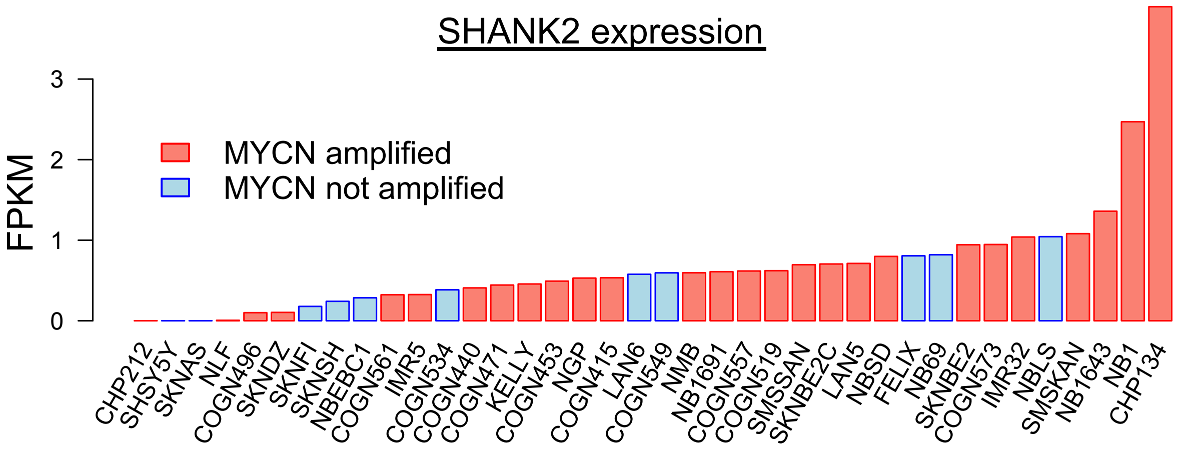

**Supplementary Figure 23: *SHANK2* expression in neuroblastoma cell lines**
*SHANK2* expression (FPKM) across 38 neuroblastoma cell lines.

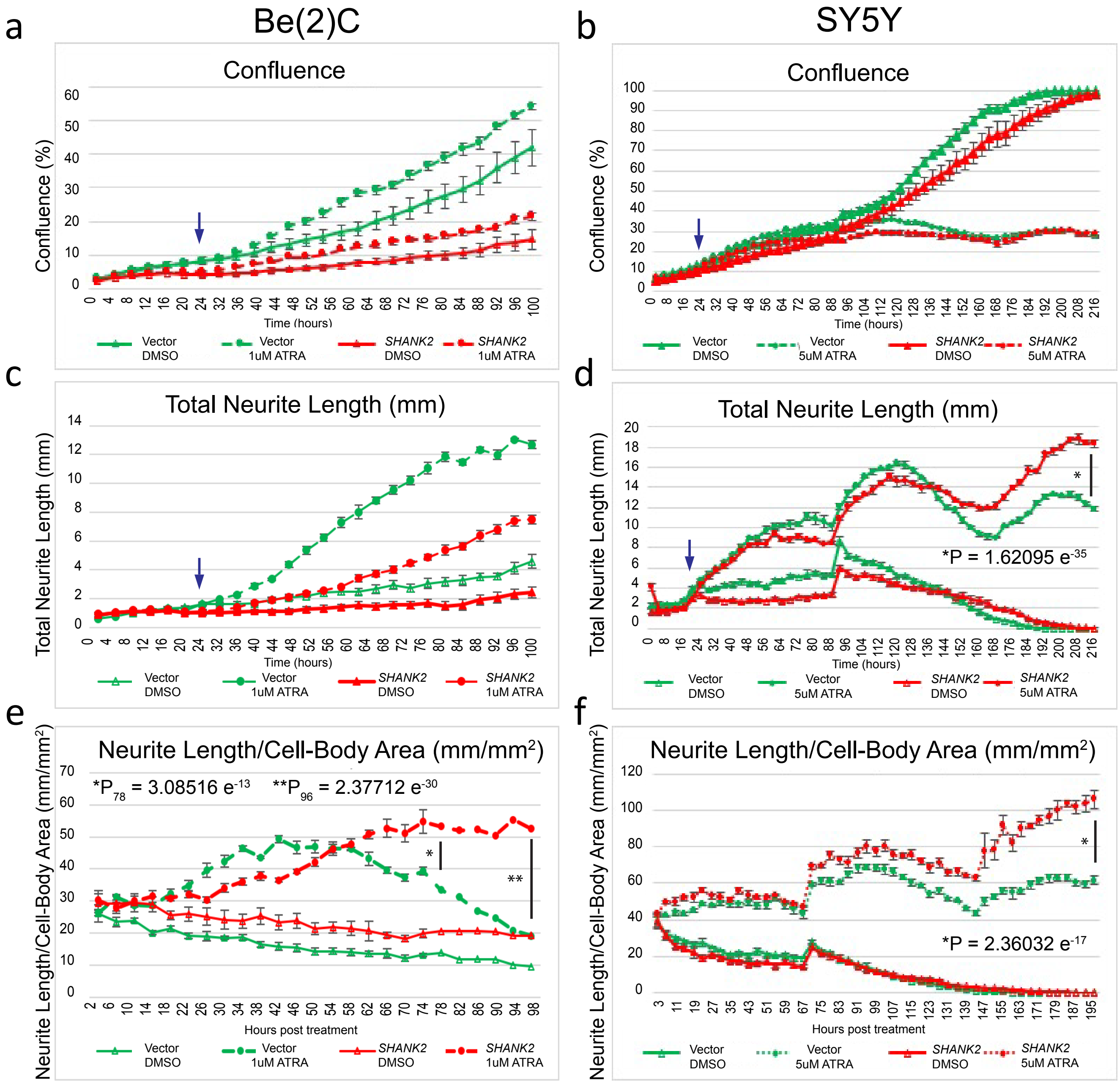

**Supplementary Figure 24: *SHANK2* accelerates differentiation of neuroblastoma cells**
**(a,b)** Confluence measures across SHANK2 expressing neuroblastoma cells, vector controls, and vehicle controls included for **(a)** Be(2)C and **(b)** SY5Y cell lines. **(c, d)** Total neurite length measurement in all cells over time for (**c**) Be(2)C and **(d)** SY5Y neuroblastoma cell lines. Arrow indicates time of ATRA introduction. **(e, f)** Neurite outgrowth normalized to cell body area over time in **(e)** Be(2)C and **(f)** SY5Y cell lines.
